# Site-Specific Crosslinking Reveals Phosphofructokinase-L Inhibition Drives Self-Assembly and Attenuation of Protein Interactions

**DOI:** 10.1101/2023.09.19.558525

**Authors:** Athira Sivadas, Eli Fritz McDonald, Sydney O. Shuster, Caitlin M. Davis, Lars Plate

**Author notes:** Corresponding author: Lars Plate. These authors contributed equally (Athira Sivadas, Eli Fritz McDonald).

## Abstract

Phosphofructokinase is the central enzyme in glycolysis and constitutes a highly regulated step. The liver isoform (PFKL) compartmentalizes during activation and inhibition *in vitro* and *in vivo* respectively. Compartmentalized PFKL is hypothesized to modulate metabolic flux consistent with its central role as the rate limiting step in glycolysis. PFKL tetramers self-assemble at two interfaces in the monomer (interface 1 and 2), yet how these interfaces contribute to PFKL compartmentalization and drive protein interactions remains unclear. Here, we used site-specific incorporation of noncanonical photocrosslinking amino acids to identify PFKL interactors at interface 1, 2, and the active site. Tandem mass tag-based quantitative interactomics reveals interface 2 as a hotspot for PFKL interactions, particularly with cytoskeletal, glycolytic, and carbohydrate derivative metabolic proteins. Furthermore, PFKL compartmentalization into puncta was observed in human cells using citrate inhibition. Puncta formation attenuated crosslinked protein-protein interactions with the cytoskeleton at interface 2. This result suggests that PFKL compartmentalization sequesters interface 2, but not interface 1, and may modulate associated protein assemblies with the cytoskeleton.

## INTRODUCTION

Metabolic diseases such as cancer (Hirschey et al., 2015) and diabetes (Guo et al., 2012) arise from cellular metabolic stress. During metabolic stress, cellular proteins form higher order structures and compartmentalization to modulate enzyme activity. For example, the yeast glycolytic enzyme glucokinase Glk1 polymerizes upon inhibition (Stoddard et al., 2020), and acetyl-CoA carboxylase polymerizes upon activation (Hunkeler et al., 2018). A glycolytic metabolon, a supercomplex of glycolytic proteins that promotes flux through glycolysis (Gavin et al., 2002), has also been hypothesized and supported based on the interactions and possible compartmentalization of different glycolytic enzymes in yeast (Kastritis and Gavin, 2018). Metabolons modulate metabolic rates through substrate channeling or sequestering of enzymatic active sites (Kastritis and Gavin, 2018; Lynch et al., 2017). Thus, higher order formation or compartmentalization of metabolic enzymes may affect the rate of metabolic processes and consequently be implicated in diseases that display dysregulated metabolism.

Glycolysis, the foundation of cellular metabolism, is rate-limited by phosphofructokinase (PFK), which converts fructose-6-phosphate into fructose 1,6 bisphosphate (Akram, 2013). Allosteric inhibitors of PFK include fructose 1,6 bisphosphate, citrate, and ATP. Allosteric activators of PFK include AMP, fructose 2,6 bisphosphate, and importantly its substrate fructose 6 phosphate (F6P) (Ha and Bhagavan, 2023; Kemp and Foe, 1983; Kemp and Gunasekera, 2002). Consequently, PFK compartmentalization into clusters and higher order structures drives many metabolic-modulating phenomena throughout biology. In C. elegans neuron, hypoxia induces the reversible formation of phosphofructokinase (PFK1.1) clusters near synapses (Jang et al., 2021). PFK1.1 interacts with glycolytic enzymes ALDO-1 (Jang et al., 2021) (aldolase) and GPD-3 (Jang et al., 2016) (GAPDH) under conditions of energy stress. Additionally, PFK-containing glycolytic metabolons have been observed in the human erythrocyte membranes (Campanella et al., 2005) and pancreatic alpha and beta cells (Ho et al., 2023). Co-clusters of PFKL, fructose 1,6 bisphosphatase, pyruvate kinase, and phosphoenolpyruvate carboxylase, which all catalyze irreversible reactions in either glycolysis or gluconeogenesis, have also been imaged in live cells and shown to direct glucose flux into the pentose phosphate pathway and the serine biosynthesis pathway, demonstrating that such compartmentalization can serve to regulate glucose-related metabolism (Kohnhorst et al., 2017).

The human liver isoform of PFK (PFKL) polymerizes into filaments upon F6P activation *in vitro* and compartmentalizes into puncta under citrate inhibition *in vivo* (Webb et al., 2017). *In vitro,* PFKL filament interactions occur at two locations: interface 1 (residues 396–401, 476–490, 510–519, and 693–705) and interface 2 (327–354, 372–377, and 712–718) (**Figure 1A**). In contrast to wild type (WT) PFKL, platelet isoform (PFKP) and a tetramer incompetent PFKL mutant F638R fail to form filaments or puncta. On the other hand, a chimeric protein comprised of the catalytic residues of PFKP (aa 1-368) and the regulatory residues of PFKL (aa 360-780) forms filaments and puncta - presumably at the same interfaces as WT as the regulatory domain contains both interface 1 and 2. Together, this suggests a relationship between activated filament formation *in vitro* and inhibitory puncta formation *in vivo* (Webb et al., 2017). However, the relationship between PFKL self-assembly arrangements and divergent protein-protein interactions that may regulate metabolic function remains unexplored. Here we explore how modulating PFKL self-assembly in cells alters PFKL protein-protein interactions with other enzymes.

**Figure 1.**
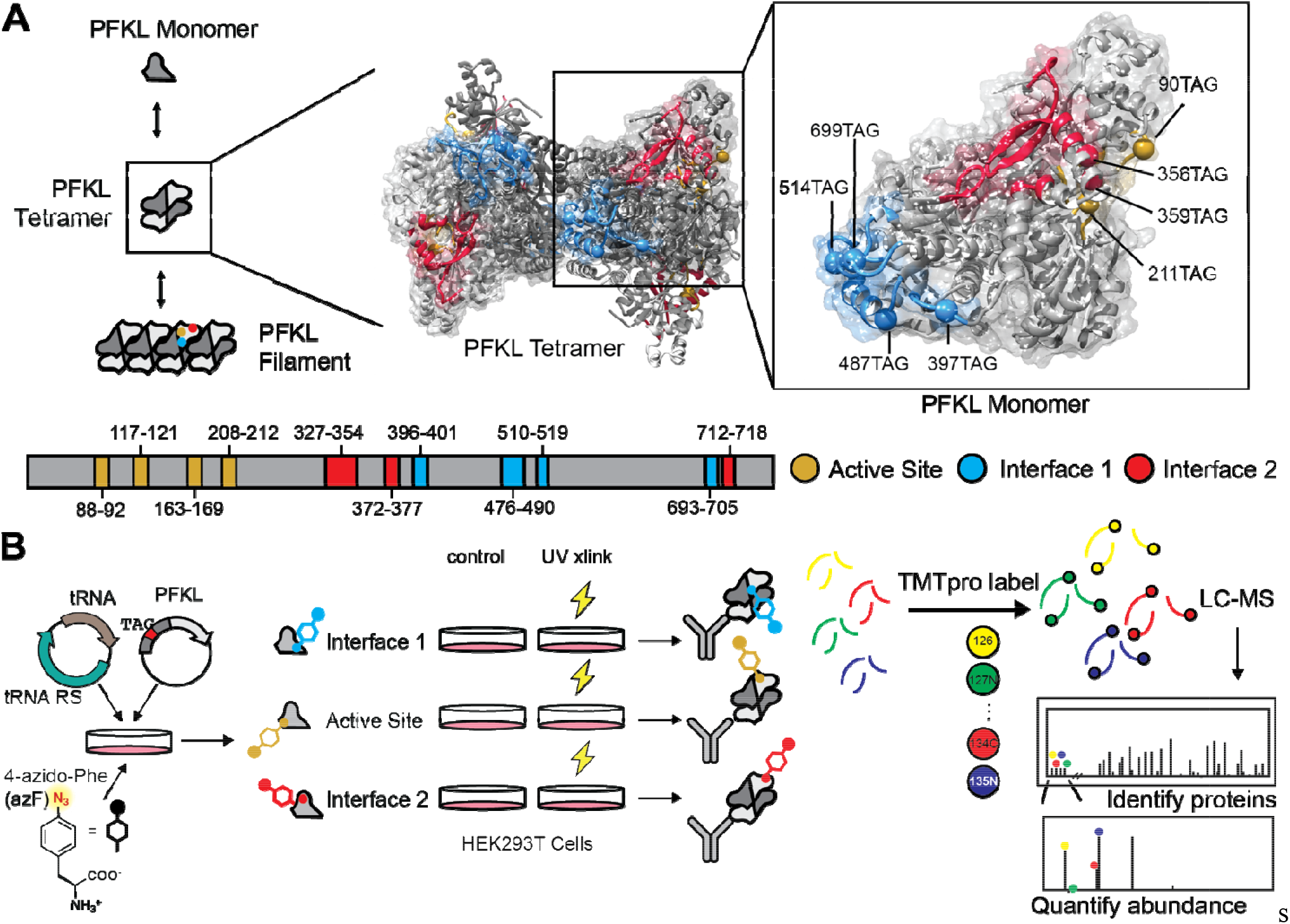
Incorporating photochemical crosslinking amino acid 4-Azido-L-Phenylalanine (AzF) site-specifically into PFKL at different filament interfaces. **A.** PFKL exists as an inactive monomer and forms tetramers to act as the third and rate-limiting step in glycolysis. PFKL also self-assembles into filaments *in vitro* and the interfaces of those filaments are shown on the tetramer and monomer for detail (PDB ID: 7LW1)(Amara et al., 2021) - the active site in gold, interface 1 s own in blue, and interface 2 shown in red(Webb et al., 2017). We introduce an amber stop codon at positions K90 and H211 in the active site, K397, Y487, Y514, and V699 in interface 1, as well as K356 and Q359 near interface 2. **B.** Schematic of the site-specific incorporation of AzF into PFKL. An AzF specific orthogonal tRNA/tRNA synthetase (RS) pair is co-transfected with PFKL containing an amber stop codon at the site of interest and cell growth media is supplemented with 1 mM AzF to allow for site-specific unnatural amino acid incorporation. Samples are then exposed to UV light to induce covalent crosslinking to interacting proteins. Finally, PFKL is immunopurified using a FLAG pulldown, crosslinked proteins are digested, TMT labeled, and pooled for LC-MS/MS quantification.

Protein-protein interactions can be quantified using affinity purification mass spectrometry (AP-MS) proteomics to reveal how structural changes in the proteome can lead to functional changes (Hashimoto et al., 2020; McDonald et al., 2022; Richards et al., 2021; Wright et al., 2021). Quantification labels such as TMT labels offer multiplexing across a variety of conditions (Li et al., 2021). However, AP-MS does not offer any insights into the binding sites for the interactors(Makowski et al., 2016). Recent advancements in chemo-proteomics resolve interactors in different protein regions by leveraging site-specific unnatural amino acid incorporation of photo-chemical crosslinking reagents to covalently crosslink interactors (Futran et al., 2015; Shah et al., 2020). However, previous studies have not quantified site-specific crosslinking interactors (Futran et al., 2015) or quantified with stable isotope labeling using amino acids in cell culture (SILAC) (Kleiner et al., 2018; Wu et al., 2020). Combining site-specific crosslinking with multiplexed labeling approaches thus offers a novel methodology for increasing throughput and characterization based on protein regions and interfaces.

Here, we used site-specific incorporation of photo-chemical crosslinking unnatural amino acid azido-phenylalanine (AzF) in combination with TMT multiplexed proteomics to investigate the dynamics and interactome of PFKL in live cells (**Figure 1B**). First, we confirmed human PFKL tolerates incorporation of AzF at several key surface exposed sites where it interfaces with filaments. UV crosslinking shifted the PFKL molecular weight observed by Western Blot, suggesting PFKL covalently crosslinked interacting proteins. Importantly, interface 1, 2, and the active site demonstrated distinct crosslinking as shown by different band shifting patterns upon UV light exposure, and interface 2 was revealed as a hub for protein-protein interactions. We further showed PFKL forms puncta under citrate-induced inhibition in human cells. Finally, we measured PFKL crosslinking using TMT labeled quantitative proteomics under citrate inhibition. Citrate inhibition decreased interface 2 protein-protein interactions with cytoskeleton components and carbohydrate secondary metabolism proteins, suggesting puncta formation sequesters interface 2 from solvent. Our examination provides insight into PFKL structure driven self-assembly and its impact on metabolism regulation.

## RESULTS

### I. PFKL tolerates substitution with AzF at active site and filament forming interfaces

To study the dynamics of PFKL self-assembly, we site-specifically incorporated the photochemical crosslinking unnatural amino acid 4-azido-phenylalanine (AzF) into an N-terminally conjugated FLAG PFKL construct (Chin et al., 2003). We substituted an amber stop codon (TAG) using site-directed mutagenesis at positions in PFKL that lie at the interfaces of PFKL filaments (Webb et al., 2017). We chose K90 and H211 near the active site, K356 and Q359 near interface 2, and K397, Y487, Y514, and V699 near interface 1 (**Figure 1A**). We refer to these PFKL mutants as K90TAG, H211TAG, etc. FLAG tagged PFKL amber stop codon constructs were transiently co-transfected with an AzF orthogonal tRNA/tRNA synthetase (Seidel et al., 2017) into HEK293T cells. Full-length PFKL mutant expression depended on the presence of 1 mM AzF in the cell growth media (**Figure 2A**), indicating the AzF was successfully incorporated into PFKL at the stop codon.

**Figure 2.**
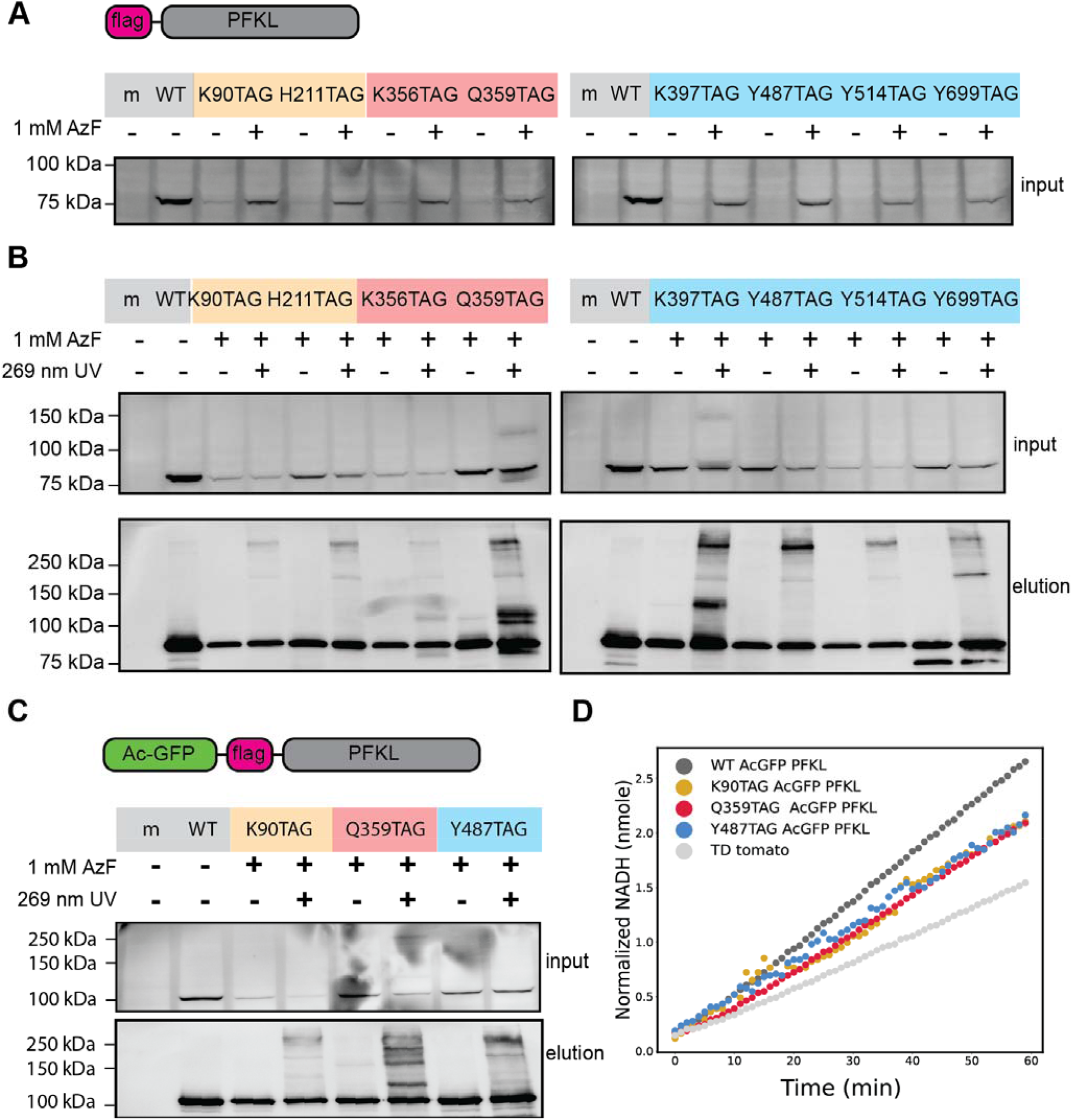
A. Representative Western blots of AzF dependent expression PFKL. AzF was incorporated into active site. interface 2 substitution mutants K90TAG, H211TAG, K356TAG, Q359TAG, and interface 1 mutants K397TAG, Y487TAG, Y514TAG, Y699TAG. In cell lysates samples (input), 85 kDa PFKL, expression depends on the presence of 1 mM AzF supplemented in the media. **B.** Representative Western blots showing 268 nm UV dependent crosslinking band shifts in PFKL active site and interface 2 substitution mutants K90TAG, H211TAG, K356TAG, and Q359TAG. Active site mutants show weak crosslinked bands at > 250 kDa. Q359TAG shows robust crosslinking bands at 100kDa, indicating an interaction with a small protein of ∼25kDa. Representative Western blots of interface 1 substitution mutants K397TAG, Y487TAG, Y514TAG, and Y699TAG. K397TAG shows robust crosslinking bands at 120kDa and > 250 kDa, Y487TAG shows crosslinking band shift at >250 kDa, and Y699TAG shows crosslinking at 150 kDa. **C.** K90TAG (active site), Q359TAG (interface 2), and Y487TAG (interface 1) show the most robust crosslinking bands and were thus selected for further analysis. Addition of the AcGFP1 fluorophore to these constructs increases the molecular weight of all bands by 25kDa. All three mutants retain the same crosslinking patterns. **D.** The activity of the different PFKL mutants with AzF incorporated was compared to that of WT PFKL for whole cell lysates. The x-axis represents the time after the reaction is initiated, and the y-axis represents [NADH] normalized to standards. [NADH] is proportional to enzyme activity. TdTomato represents a transfection control with only endogenous PFKL, while all other samples have endogenous PFKL as well as transfected PFKL.

### II. PFKL has unique crosslinking patterns at the active site and interfaces

*In vitro,* PFKL self assembles into filaments along two interfaces (**Figure 1A**)(Webb et al., 2017). We sought to measure PFKL interactions at the resolution of each interface using site-specific crosslinking *in vivo*. AzF-incorporated PFKL constructs were exposed to 269 nm UV light for 1.5min to determine if PFKL would covalently crosslink interacting molecules in an interface dependent manner. PFKL band shift to a higher molecular weight would indicate covalent crosslinking (Kleiner et al., 2018; Wu et al., 2020). Indeed, we observed the Q359AzF PFKL band shift from 85 kDa to 110kDa, the K397AzF band shift from 85 kDa to 120 kDa, and the 487TAG band shift 250kDa upon UV light exposure (**Figure 2B**). By contrast, 90TAG and 211TAG failed to show strong band shift although a faint band is seen at >250 kDa. This suggests the active site crosslinks few or no interacting proteins (**Figure 2B**). Alternative sites K356TAG in interface 2 as well as Y514TAG and V699TAG in interface 1 demonstrated similar albeit less robust band shifting (**Figure 2B**). Thus, we proceeded with 359TAG and 487TAG for quantitative mass spectrometry to investigate interactions with interface 2 and 1 respectively and included 90TAG in the active site as a control.

To visualize PFKL self-assembly using fluorescence microscopy, we added an AcGFP1 tag N-terminally to the FLAG tag. AcGFP1 tagged PFKL constructs K90TAG, Q359TAG, and Y487TAG were crosslinked to ensure reproducible band shift observed as for FLAG-tagged PFKL. Of note, AcGFP1 added 23 kDa to PFKL, making the WT band appear at 110 kDa. We observed faint band shifting in K90TAG, robust band shifting in Q359TAG from 110 kDa to 135 kDa, and robust Y487TAG band shifting from 100 kDa to 250 kDa (**Figure 2C**) with the addition of AcGFP1. Reproducing the band shifting patterns with the AcGFP1 tag demonstrated the fluorophore did not alter crosslinking.

PFKL filament formation requires maintenance of enzymatic function (Webb et al., 2017). To ensure biological relevance of our methodologies, we determined that AzF-substituted PFKL constructs maintained enzymatic function, by performing a PFK activity assay. When compared to TD tomato transfected HEK293T controls, WT PFKL expressing cell lysates showed an increase in PFK activity as measured by NADH production (**Figure 2D**). Furthermore, 90TAG, 359TAG, and 487TAG PFKL transfected HEK293T cell lysates demonstrated PFKL activity above TD tomato controls but below WT (**Figure 2D**). This confirmed mutant activity above mock transfected lysate, but also suggest amber stop codon substitution affects expression or activity directly compared to WT.

### III. PFKL interface 2 crosslinks robustly with cytoskeletal proteins

To investigate the proteins involved in differential band shifting patterns observed across the PFKL interfaces, we performed multiplexed TMT labeled affinity purification mass spectrometry (AP-MS) to quantitatively compare protein levels between interface 1 and 2 and the active site (**Figure 1B**). (Kim et al., 2023; McDonald et al., 2022; Plate et al., 2019; Wright et al., 2021). Our results suggest that crosslinking changes protein levels measured after AzF-incorporation at interface 2 but not the active site or interface 1.

We transiently co-transfected tdTomato, WT, K90TAG, Q359TAG, and Y487TAG PFKL with AzF tRNA/RS into HEK293T cells and supplied 1 mM AzF supplemented media for at least 24 hours. AzF substituted constructs were exposed to 269 nm UV light and non-exposed controls were also collected. PFKL was co-immunoprecipitated (Co-IP) from cell lysates and washed 4 times in radio-immunoprecipitation assay (RIPA) buffer to stringently remove non-crosslinked proteins (Chen and Peng, 2020). The protein elution samples were digested into peptides with trypsin, TMT pro 18plex regents were used to label individual samples (**Supplemental Table 1**), and then samples were pooled and analyzed by multidimensional protein identification technology (MudPIT)-tandem MS(Li et al., 2021).

Peptides were identified from tandem mass spectra, and importantly TMT reporter ions were produced during peptide fragmentation that enabled quantitative, relative comparison of peptide abundances across the different Co-IP conditions (**Figure 1B**). We conducted Co-IPs and independent MS runs for multiple replicates per mutation (WT *N* = 3, K90AzF *N* = 4, Q359Azf *N* = 4, Y487AzF *N* = 4) (**Supplemental Table 1**). To distinguish PFKL-specific interactors, we included a TdTomato mock transfection control to account for nonspecific interactors that have high affinity for the bead matrix during the Co-IP. Statistically significant interactors were filtered by considering the log2 TMT intensities of protein levels in the PFKL Co-IP samples over the tdTomato mock transfected samples. To compare TMT quantification, we used median normalized TMT intensity based on the assumption that the mock and PFKL/FLG-AcGFP1-bait Co-IP samples contain similar levels of nonspecific background proteins (**Supplemental Figure S1A-B**). Each statistically significant interactor was gathered in the master list of interacting proteins across all sample conditions (**Supplemental Table 2**).

The volcano plot for WT PFKL confirmed efficient pull down of the PFKL bait protein (divided into two different peptide sequences to separately quantify the PFKL FLAG-AcGFP1 tag) (**Figure 3A**). FLAG-AcGFP1 is comprised of the sequence of FLAG, thrombin, 6xHis, and AcGFP1 at the N-terminus of the PFKL constructs. PFKL sequence by contrast quantifies both transiently transfected PFKL as well as endogenous PFKL. Notably, the stringent bead washes successfully removed most non-covalent interacting proteins as demonstrated by the lack of high confidence proteins besides the baits (**Figure 3A**).

**Figure 3.**
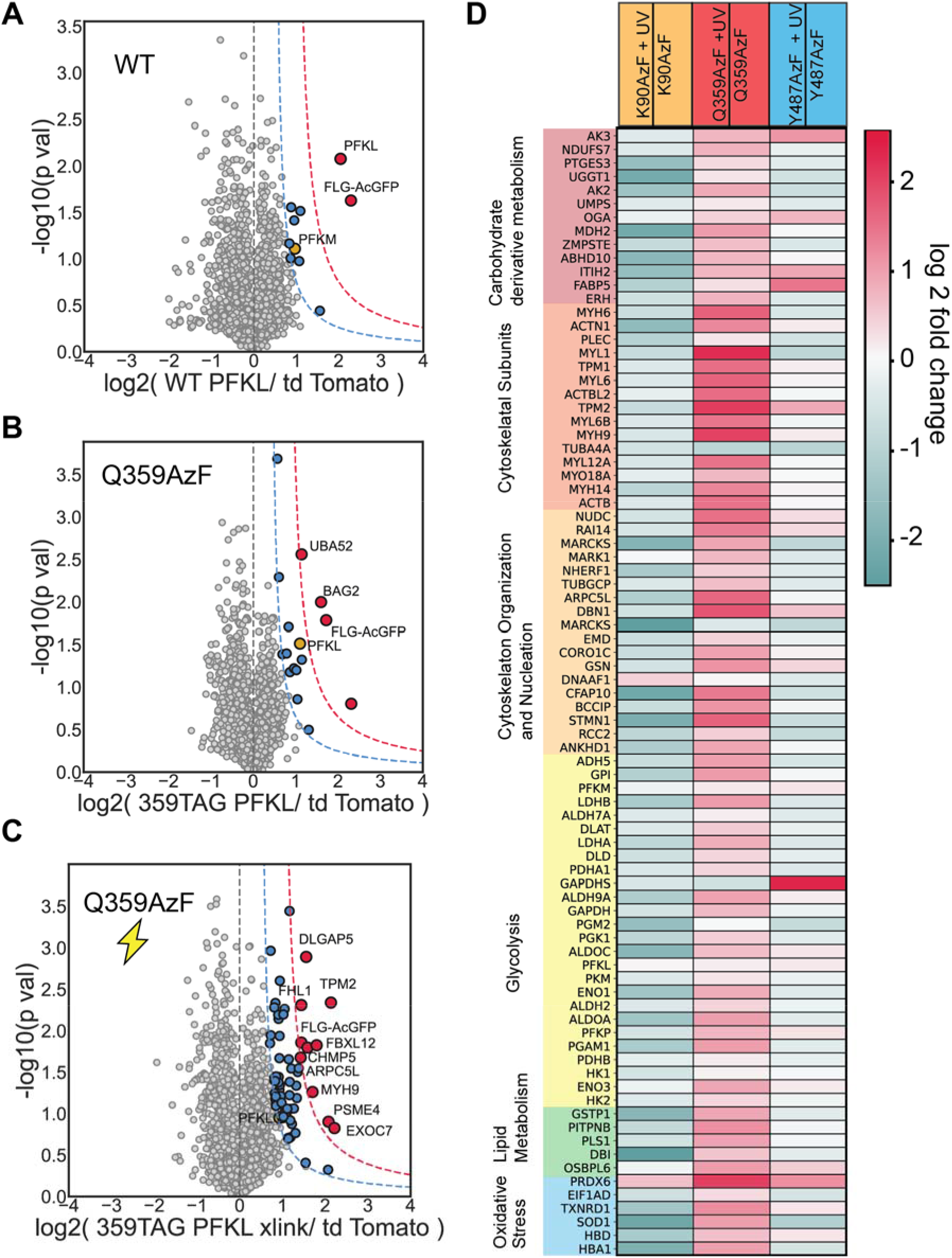
Covalent crosslinking successfully enriches interacting proteins at interface 2 as measured by Q359AzF. **A.** Volcano plot of WT PFKL over tdTomato mock transfection control for identification of interactors. The x-axis represents the log2 fold change of WT PFKL TMT quantification over the tdTomato TMT quantification across three biological replicates. The y-axis represents the −log10 transformed p-values, as calculated by a paired student’s t-test. Bait protein is labeled as PFKL and FLG-AcGFP1. Enrichment of bait proteins and lack of interactors demonstrates stringent washing steps remove non-covalent interacting proteins. High-confidence (red) and medium-confidence (blue) interactors are demarcated by curve cut off at 1σ’ and 2σ’ of the log2 fold change distribution respectively. Glycolytic interactors are represented in yellow. **B.** Volcano plot of Q359AzF over tdTomato mock transfection control for identification of interactors. Colored same as A (n = 4 biological replicates). **C.** Volcano plot of Q359AzF exposed to UV light and hence covalently crosslinked to interacting proteins compared tdTomato mock transfection control interactors across (n = 4). An increase of upregulated, statistically significant interactors indicates crosslinking was successful. Colored same as A. **D.** Heatmap comparing enrichment of interactors functioning in different cellular pathways (categorized according to GO-terms biological process) among the three mutants. The interactors enriched in the crosslinked samples are normalized to those enriched in the non-crosslinked samples. The log2 fold change is color coded (teal low, pink high enrichment). Site Q359 of PFKL near interface 2 demonstrates robust interactions with cytoskeletal subunits and a higher number of interactions overall compared to the other two sites.

Likewise, the volcano plot for interface 2 substitution Q359AzF showed a lack of high confidence interactors indicating stringent washes removed most non-covalent interactions (**Figure 3B**). Interestingly, the volcano plots for Q359AzF + UV conditions demonstrated a clear increase in the number of medium and high confidence interactors (**Figure 3C**) indicating that AzF crosslinking increases the abundance and confidence of interacting proteins. By contrast, volcano plots of active site substitution K90AzF and interface 1 substitution K487AzF showed similar levels of medium and high confidence interactors with and without crosslinking (**Supplemental Figure S1C-F**). Together, these data implicated interface 2 as a hotspot of PFKL interactions *in situ*.

We sought to understand the biological significances of interface 2 interactions; thus, we annotated each identified interactor according to biological process GO-term (**Supplemental Table 3**). We plotted a heatmap of the log2 fold change of each crosslinked conditions (position 90, 359, and 487) over their respective non-crosslinked controls (**Figure 3D**). Pathways carbohydrate derivative metabolism, cytoskeleton organization and subunit, glycolysis, lipid metabolism, and oxidative stress were selected for their presumed role in glycolysis and adjacent pathways (Amara et al., 2021; Park et al., 2020). Notably, pathways important for PFKL, such as cytoskeleton and glycolysis, have a distribution of higher log2-fold change over mock controls when compared with pathways less clearly associated with PFKL such as DNA, RNA, Vesicular Transport, and Signaling (**Supplemental Figure S2A-L**).

Compared to K90TAG and Y487TAG, K359TAG demonstrated more robust interactions in all five pathways. For instance, in regard to the carbohydrate derivative metabolism pathway, extensive interaction with malate dehydrogenase, an enzyme of the Krebs cycle, was observed. With regards to the cytoskeletal pathway, extensive interaction with actin, actin-associated proteins, and proteins that regulate actin, such as ACTB, MYL1, DBN1, and ARPC51 was observed. As for the glycolysis pathway, interface 2 seemed to interact quite robustly with GPI, or glucose-6-phosphate isomerase, which catalyzes the step preceding the PFK reaction. Interaction with ALDOA, aldolase A, which catalyzes the step after the PFK reaction, was also observed. Interestingly, within the lipid metabolism pathway, interaction with PITPNB, a transferase of membrane lipids, was observed, and at the intersection of lipid metabolism and oxidative stress, strong interaction with PRDX6, an enzyme with phospholipase activity, was observed. These data further indicated that interface 2 crosslinks at greater intensity and to a more diverse set of partner proteins compared to the active site and interface 1 (**Figure 3D**).

### IV. Citrate treatment induces PFKL puncta in WT and AzF substituted PFKL mutants

Citrate treatment leads to PFKL puncta formation in rat cells (Webb et al., 2017) and human cells (Kohnhorst et al., 2017). We sought to determine if PFKL formed puncta in our transiently expressing HEK293T system was amenable to amber stop codon suppression technology. Briefly, we transfected WT AcGFP1-flag-PFKL DNA into HEK293T cells, waited approximately 16-18 hours, and added 20 mM of sodium citrate (pH to 7.5) to the media. WT AcGFP1-PFKL indeed exhibited localization into small clusters of increased intensity compared to diffuse GFP fluorescence in control conditions (**Figure 4A-B**), highlighting that puncta formation occurred due to citrate-induced PFKL inhibition in human cells.

**Figure 4.**
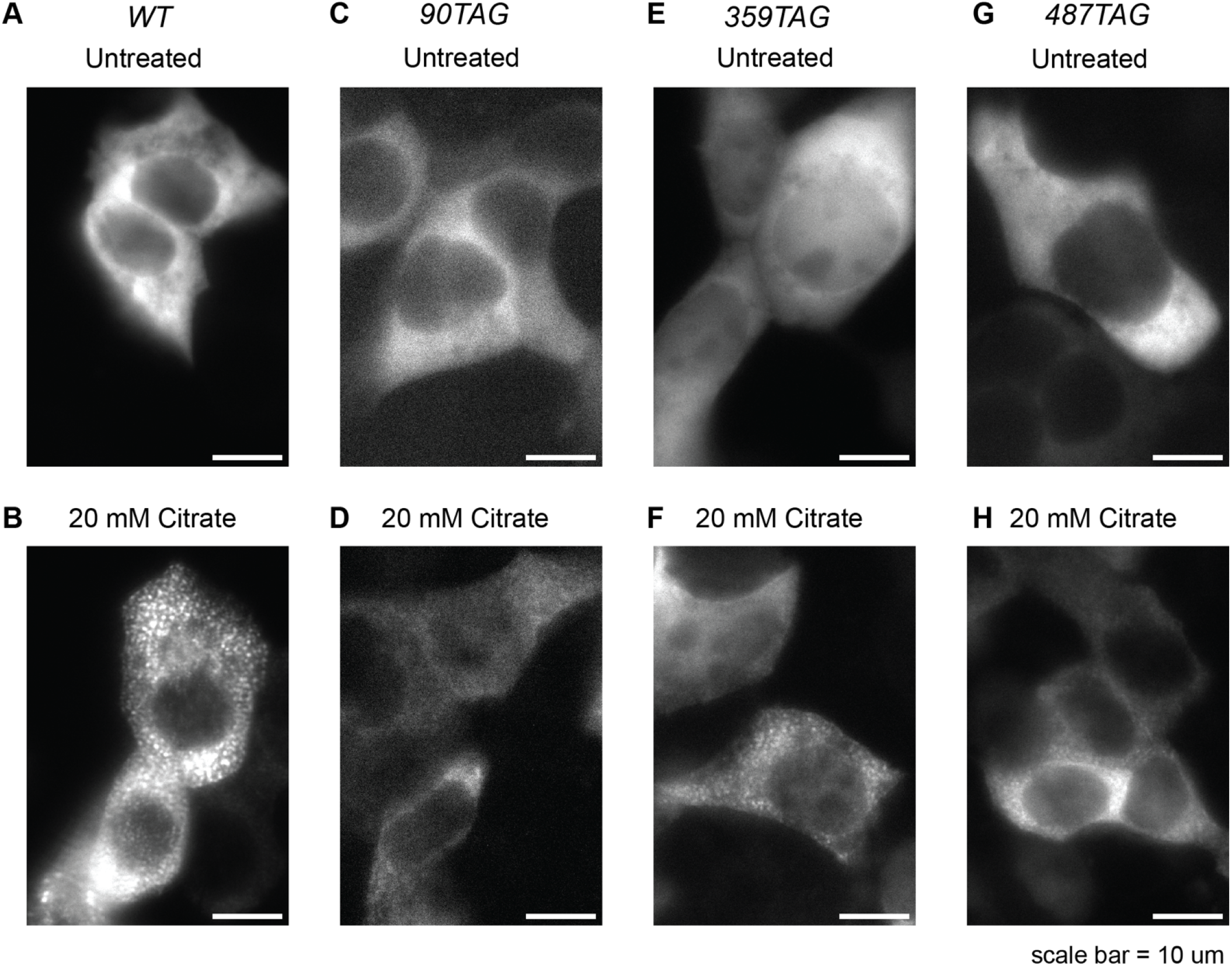
Confocal microscopy of HEK293T cells transiently transfected with AcGFP1-PFKL. Experimental samples were imaged 10 minutes after addition of citrate. **A.** WT PFKL in basal conditions. Fluorescence is diffuse, indicating that PFKL is dispersed throughout the cell. **B.** WT PFKL after addition of 20mM citrate. The fluorescence forms localizations around the periphery of the cell, indicating that PFKL is sequestering into puncta. **C.** 90TAG AcGFP1 PFKL mutant in basal conditions. **D.** 90TAG AcGFP1 PFKL mutant after addition of 20mM citrate. Localizations of fluorescence are still observed, but to a lower extent than seen for WT PFKL. **E.** 359TAG AcGFP1 PFKL mutant in basal conditions. **F.** 359TAG AcGFP1 PFKL mutant after addition of 20mM citrate. **G.** 487TAG AcGFP1 PFKL in basal conditions. **H.** 487TAG AcGFP1 PFKL after addition of 20mM citrate.

Next, we examined the localization dynamics of active site AzF-substituted PFKL under conditions of citrate inhibition. We transiently co-transfected K90TAG FLAG-AcGFP1-PFKL and AzF tRNA/RS DNA into HEK293T cells, added 1 mM AzF supplemented media 16 hours later and after 2 hours switched to Fluorobrite media supplemented with 10 % FBS and imaged the AcGFP1 fluorescence on an inverted epifluorescent microscope before and after addition of 20 mM sodium citrate (pH to 7.5 with NaOH). We observed a primarily diffuse GFP signal from the treated K90AzF FLAG-AcGFP1-PFKL compared to untreated control with limited signs of puncta formation (**Figure 4C-D**). This was in line with K90AzF FLAG-AcGFP1-PFKL showing lowest expression. We suspect that some free AcGFP1 expression due to premature stop codon truncation may obscure the puncta-localized fluorescence. However, consistent with previous findings associated with puncta (Webb et al., 2017), we still observed PFKL migration and compartmentalization in the cell periphery of the cell after addition of citrate.

Finally, we examined the citrate inhibition reaction of interface 1 and 2 AzF substituted Y487TAG-FLAG-AcGFP1-PFKL and Q359TAG-FLAG-AcGFP1-PFKL respectively. Again, we transiently transfected these respective constructs with AzF tRNA/RS DNA into HEK293T cells, added 1 mM AzF supplemented media, and took fluorescence images before and after addition of 20 mM sodium citrate (pH to 7.5). Representative images show citrate induced some puncta in AzF substituted interface 2 Q359AzF (**Figure 4E-F**) and puncta and peripherally localized PFKL in interface 1 Y487AzF (**Figure 4G-H**). Together, these data suggest AzF substitution in PFKL interfaces retains self-assembling and clustering physiology in our expression system. Thus, we next sought to measure the effect of citrate PFKL protein crosslinking with quantitative proteomics to investigate how puncta formation alters the engagement of protein interactions at the respective PFKL assembly interfaces.

### V. Citrate induced puncta drives crosslinking changes at PFKL interface 2

Citrate treatment causes PFKL mutants with AzF to self-assemble into puncta. We sought to measure changes in site-specific crosslinking under citrate conditions to test if puncta sequestered or exposed specific interfaces of PFKL.

TdTomato (mock), WT, Y90TAG, Q359TAG, and Y487TAG PFKL were transiently co-transfected with AzF tRNA/RS into HEK293T cells with 1 mM AzF. To prevent excessive cell stress and death, 10 mM citrate was added for 60 seconds and an additional 10 mM citrate for 60 seconds, bringing total concentration to 20 mM. Briefly, UV exposed and unexposed plates were collected, immunoprecipitated, washed, eluted, chloroform methanol precipitated and digested by trypsin into constituent peptides. We then labeled each sample with TMT pro 18plex regents and pooled samples for mass spectrometry (Li et al., 2021). We considered the log2 TMT intensities of protein levels in the PFKL Co-IP samples over the tdTomato mock transfected samples to determine statistically significant interactors.

Since interface 2 had robust changes in interactor abundance upon UV exposure (**Figure 3B-D**) we measured these crosslinking changes in the presence of 20 mM citrate. Under citrate treatment, Q359AzF volcano plots showed successful Co-IP of PFKL bait as quantified by PFKL and FLAG-AcGFP1 abundance (**Figure 5A**). However, the presence of several high confidence interactors suggests the pull down was not as clean as Q359AzF without citrate. When we crosslinked Q359AzF +20 mM citrate, we observe a decrease in the number of high confidence interactors overall, indicating that citrate-induced puncta formation sequestered interface 2 of PFKL (**Figure 5B**). Active site and interface 1 volcano plots showed similar levels of medium and high confidence interactors regardless of crosslinking (**Supplemental Figure S3A-D**).

**Figure 5.**
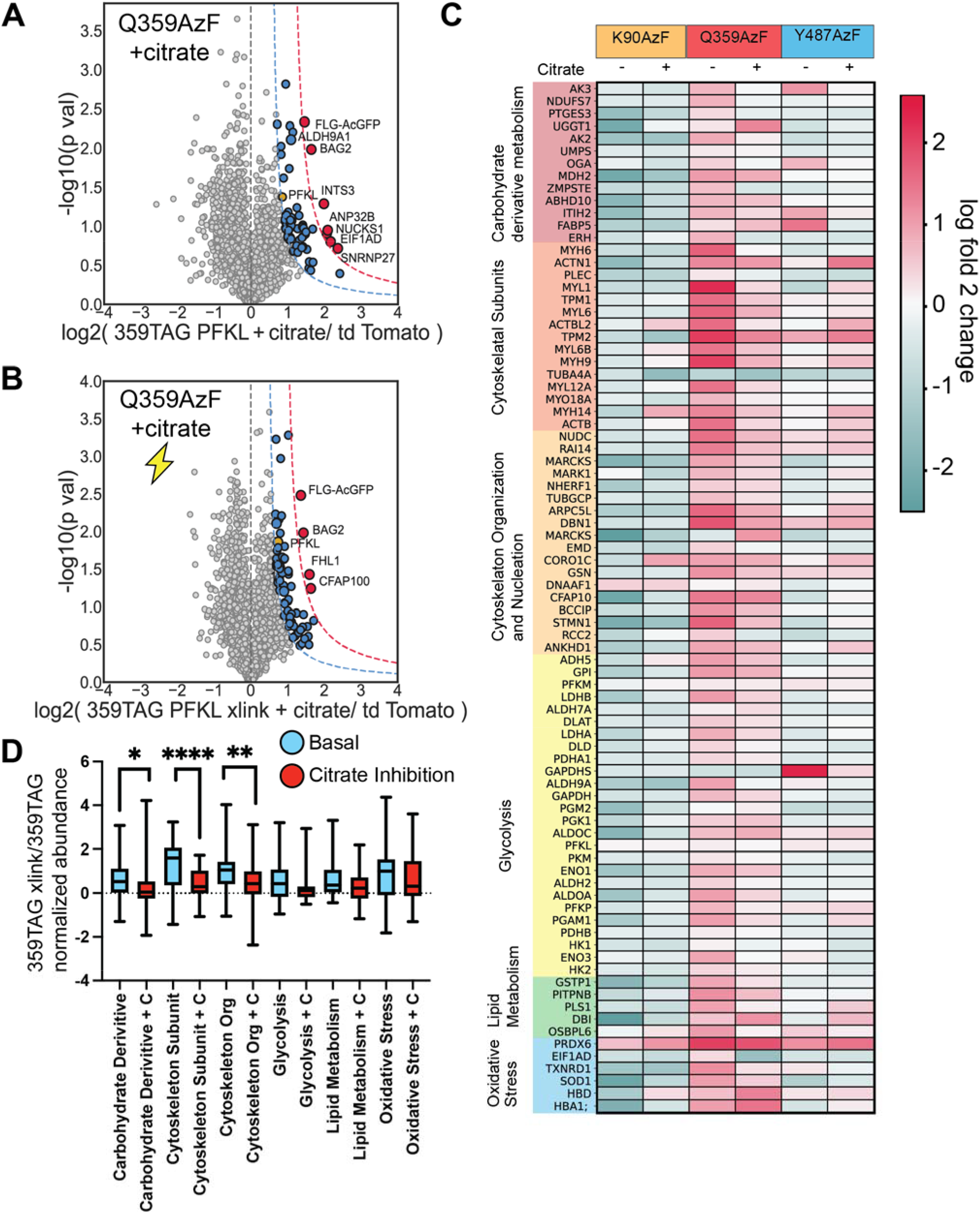
Sodium citrate treatment attenuates interaction with cytoskeletal and carbohydrate derivative metabolism proteins at interface 2. **A.** Volcano plots of Q359AzF with 20 mM citrate. The x-axis represents the log2 fold change of interactors between Q359AzF treated citrate compared to the tdTomato control (n = 4 biological replicates). The Y axis represents the −log10 transformed p-values, as calculated by student’s t-test. Bait protein is labeled as PFKL and FLG-AcGFP1. High-confidence (red) and medium-confidence (blue) interactors are demarcated at the curve cut offs over 1σ’ and 2 σ’ respectively. Glycolytic interactors are represented in yellow. **B.** Volcano plots of Q359AzF with 20 mM citrate exposed to UV light across. Colored the same as A (n = 4). **C.** Heatmap comparing enrichment of interactors functioning in different cellular pathways (categorized according to GO-terms biological process) among the three assessed mutants. Each row represents the log2 fold change of crosslinked samples (e.g. + UV light) over the respective sample without crosslinking (e.g. no UV light). The addition of citrate to both the crosslinked and non-crosslinked samples in indicated. The interactors enriched in the crosslinked samples are normalized to those enriched in the non-crosslinked samples. The log2 fold change is color coded (teal low, pink high enrichment). **D.** Aggregated interactions by pathways from **C.** in the 359TAG xlink/359TAG and 359TAG xlink + citrate /359TAG + citrate conditions.

To understand the biological significance of interface 2 interactions, we plotted a heatmap containing our master list of protein interactions (**Supplemental Table 2**) of the log2 fold change of crosslinked conditions over non-crosslinked controls with citrate treatment (**Figure 5C**). We again examined the upregulation of cytoskeleton and glycolysis proteins against unrelated pathways DNA, RNA, Signaling, and Vesicular Transport, and Signaling. We found relevant pathways have a higher log2-fold change distribution of compared unrelated pathways (**Supplemental Figure S4A-L**). These data suggest that during citrate PFKL inhibition interface 2 has become sequestered and consequently shows attenuated crosslinking and enrichment of interactors (**Figure 5C**).

This observation is consistent with cryo-EM structures of PFKL activated filaments where interface 2 was sequestered by interface 1 of another tetramer of PFKL (Webb et al., 2017). PFKL itself may have physically obstructed interface 2 during puncta formation, preventing interactors from crosslinking. However, since interface 1 crosslinked interactors changed little during inhibition, it remains unclear if interface 2 is sequestered by precisely the same interface arrangement as shown *in vitro* activated filaments.

To gain insights into how citrate inhibition impacted specific pathways interactions at interface 2, we aggregated the quantified interaction changes for each pathway (**Figure 5D, Supplemental Table 4**). Citrate significantly reduced interactions at interface 2 with carbohydrate derivative metabolism, cytoskeletal subunit proteins and cytoskeletal organization/nucleation proteins (**Figure 5D**). These data suggested citrate induced puncta sequestered interface 2, preventing it from crosslinking to interactors.

### VI. Cytoskeleton Organization Proteins Link Metabolism Proteins to the Cytoskeleton

Previous research suggested glycolysis–cytoskeleton interactions played a role in regulating cell response to changes in metabolic flux (Masters, 1984; Park et al., 2020; Real-Hohn et al., 2010). Citrate inhibition statistically significantly decreased PFKL-cytoskeleton interactions, as measured by site specific crosslinking at interface 2, across two cytoskeletal pathways: cytoskeletal subunit proteins and cytoskeletal organization/nucleation proteins (**Figure 5D**). Additionally, citrate inhibition decreased interface 2 interactions with carbohydrate derivative metabolism proteins (**Figure 5D**). However, the spatial connections between these pathways: glycolysis, carbohydrate derivative metabolism, and cytoskeletal proteins, remains unclear.

To empirically map the relationship between these pathways we created a network plot of protein-protein interactions from our master list of interactors by conducting a secondary extended interactome search on the STRING database (Szklarczyk et al., 2019). Briefly, primary PFKL interactors were searched for their top 20 interactors on STRING to create a data set of secondary extended interactors. Then all secondary interactors were cross referenced against each other, and non-overlapping secondary interactors were dropped from the dataset. This previously established protocol (Davies et al., 2020) creates a network of closely interacting proteins that reveals clusters of protein pathways in metabolism, nucleobase biosynthesis, and cytoskeleton proteins (**Figure 6**). Importantly, CTNNB1, RCC2, AURKB, NME7, TP53, and STMN1, proteins involved in cell cycle, linked metabolic proteins to cytoskeletal proteins among our extended interactome network (**Figure 6**). Cell cycle proteins implies a role in nucleating polymerization of cytoskeletal subunits as occurs during mitotic spindle formation and cell division. Notably few secondary interactors overlapped between glycolytic and cytoskeletal subunit proteins, implying changes observed with citrate inhibition may be mediated through cytoskeletal organization proteins instead. These data validated our crosslinking approach as glycolytic metabolons formation near the cytoskeleton have been demonstrated (DeWane et al., 2021; Kastritis and Gavin, 2018; Masters, 1995, 1984; Norris et al., 2013; Park et al., 2020; Stoddard et al., 2020). Additionally, opposed to cytoskeletal subunit proteins, our results suggest a previously underappreciated role of cytoskeleton reorganization during cell division proteins during changes in metabolic flux.

**Figure 6.**
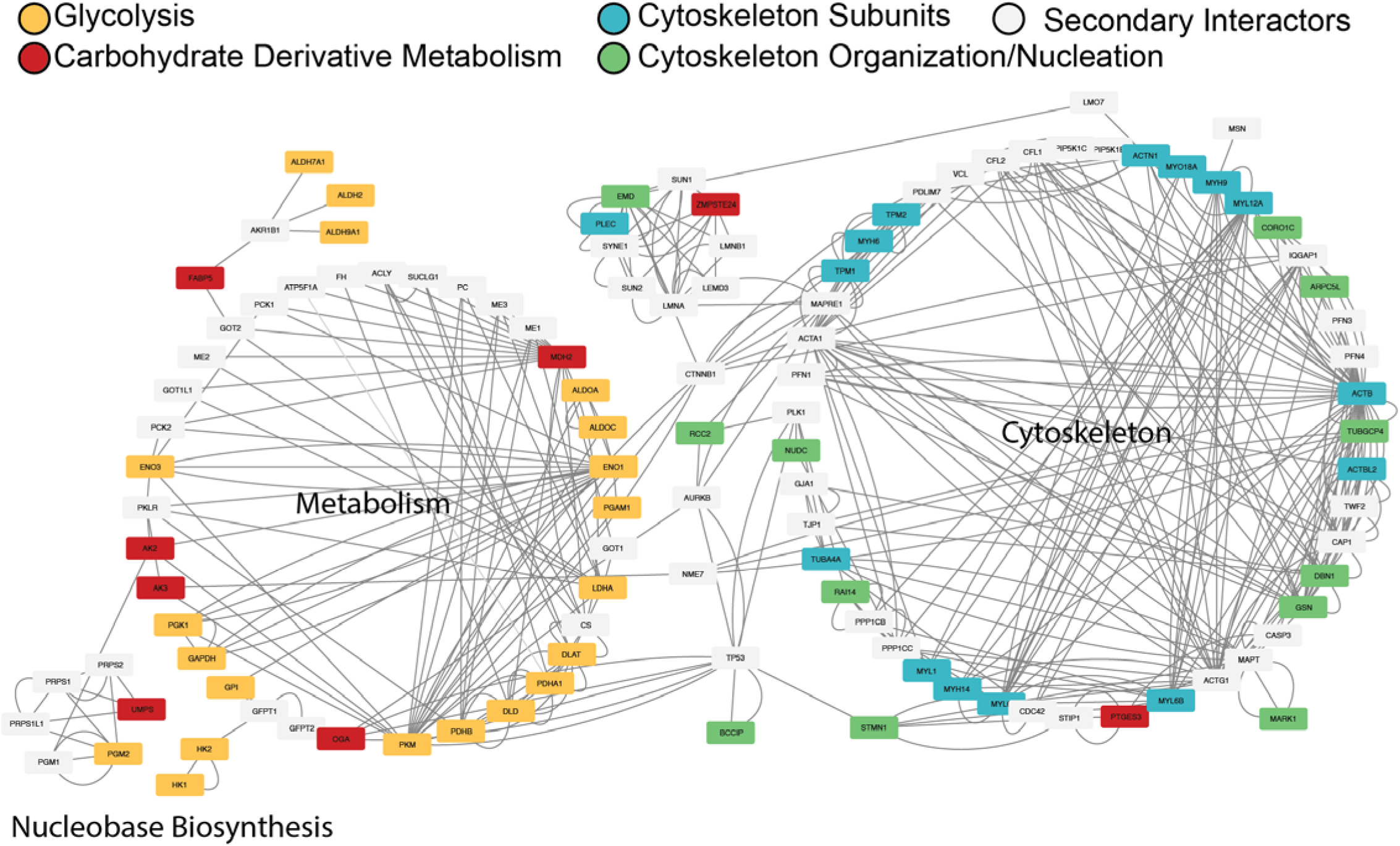
Protein interaction network reveals clusters between identified PFKL interactors. STRING database search was performed for the top 20 interacting proteins. Extended and overlapping interactors between identified PFKL interactions in four important pathways: Glycolysis, Carbohydrate Derivative Metabolism, Cytoskeleton Subunit proteins, and Cytoskeleton Organization/Nucleation proteins. Primary PFKL interactors in the Glycolysis pathway are shown as gold rectangles, primary PFKL interactors in the Carbohydrate Derivative Metabolism are shown as red rectangles, primary PFKL interactors with Cytoskeleton Subunit proteins are shown as blue rectangles, and primary PFKL interactors with Cytoskeleton Organization proteins are shown as green rectangles. Overlapping secondary interactors scraped from the STRING database are shown as grey rectangles. PFKL, which interacts directly with all colored proteins, and proteins that failed to demonstrate any overlapping interactors were excluded for clarity.

## DISCUSSION

PFKL distinguishes itself from other human PFK isoforms by its ability to compartmentalize at specific interfaces during metabolism modulation (**Figure 1A**). However, the impact of PFKL compartmentalization on other metabolic enzymes and cellular components remains poorly understood. To investigate PFKL self-assembly impact on the cell we measured interactors using site-specific incorporation of photochemical crosslinking amino acids. We verified that PFKL tolerates incorporation in interface 1, interface 2, and the active site and showed upregulation of interface 2 interactors with the cytoskeleton in the presence of UV light.

We observed puncta formation in human embryonic kidney cells under conditions of citrate inhibition and examined how this structural change altered the PFKL interactome. Although site 359 (interface 2), site 487 (interface 1), and site 90 (active site) all demonstrated band shifting by Western blot via UV-induced crosslinking, site 359 (interface 2) showed the most robust crosslinked interactions by mass spectrometry, as well as changes in interactome under citrate conditions.

Solvent-accessible interfaces of the protein can crosslink interactors. During citrate-induced puncta formation, we hypothesize interface 2 is sequestered from the solvent and therefore reduces the log2 fold change of its interactors. Most notably, reductions in interactions are observed in the pathways carbohydrate derivative metabolism and cytoskeleton subunits, as well as cytoskeleton organization and nucleation. By contrast, global interactions under citrate conditions versus basal conditions showed less prominence at interface 1 and the active site.

PFK has been extensively shown to interact with other enzymes of glycolysis and glucose metabolism. We provided evidence that interface 2 interacted with many of these enzymes, such as aldolase A and phosphoglycerate mutase, and these interactions were reduced, albeit not significantly, in the presence of the PFKL allosteric inhibitor citrate. Formation of a glycolytic metabolon and interactions between glycolytic enzymes (Kastritis and Gavin, 2018) increased the activity of these enzymes (Campanella et al., 2005; Ho et al., 2023; Menard et al., 2014). Thus, we expected the opposite to also hold: PFKL inhibition decreases interactions with glycolytic proteins. Citrate-induced puncta formation of PFKL may serve to reduce flux through glycolysis by sequestering PFKL into compartments and removing the enzyme from the proximity of other glycolytic enzymes. These structural changes likely impaired glycolytic metabolon formation.

F6P activated PFKL filaments *in vitro* assemble through interactions between interface 1 and 2 (Webb et al., 2017). Citrate-induced puncta formation may sequester interface 2 to prevent assembly that enhances enzyme activity. Notably, citrate binds the PFK C-terminus, evident by attenuated inhibition in PFKM mutants K557R or K617A (Usenik and Legiša, 2010). Binding proximate to interface 1, albeit not overlapping, may prevent interface 1 from participating in self-assembly with interface 2 during citrate inhibition – leading to a distinct arrangement of filamentous PFKL found in puncta. This suggests a possible relationship for the opposing self-assembly structures observed between activation and inhibition.

Interface 2 of PFKL also demonstrated a similar trend of crosslinking to cytoskeletal subunits. Interactions with these proteins decreased significantly during citrate inhibition. This finding has implications in the cytoskeletal integrative sensor hypothesis, which proposes that the binding of cytoskeletal proteins to metabolic enzymes allows for the regulation and integration of cellular metabolism (Norris et al., 2013). This hypothesis suggests that cytoskeletal binding of enzymes may impact their activity (Masters, 1984). Elevated rates of glycolysis have been shown to increase binding of glycolytic enzymes to the cytoskeleton (Masters, 1984; Masters and Wilson, 1981). Binding of glycolytic enzymes to f-actin is stimulatory (El-Bacha et al., 2003; Gomes Alves and Sola-Penna, 2003; Masters, 1984), whereas binding to microtubules is inhibitory (Ovádi et al., 2004; Ovádi and Saks, 2004). We observed PFKL interactions with actin and actin-associated proteins, but comparatively little interaction with microtubules (**Figure 3D & 5C).**

Actin may even serve as a scaffold for the formation of a glycolytic metabolon (Masters, 1995). In erythrocytes, actin association with PFK1 has been shown to stimulate enzyme activity and actin disassociation from PFK1 has been shown to enhance enzyme inhibition^16^. Specifically, actin binding stabilizes the tetrameric form of PFKL or its fully active form, lowers its K_M_, and reduces the enzyme’s susceptibility to allosteric inhibitors such as ATP, citrate, and lactate (DeWane et al., 2021). These findings were consistent with our observations that PFKL interacted with actin-associated proteins less during conditions of citrate-induced inhibition. As with the glycolytic proteins, the citrate-induced puncta could have a sequestering effect on PFKL that physically distances the enzyme from actin to suppress enzyme activity.

The method of site-specific unnatural amino acid incorporation has been traditionally used to fluorescently label proteins, enhance antibody-drug interactions, investigate protein structure and function, etc. (Zhang et al., 2013). We present a novel approach of investigating changes in site-specific interactions by integrating unnatural amino acid incorporation experiments with multiplexed TMT labeled mass spectrometry proteomics. Such a technique has allowed us to not only identify the interactors of PFKL at specific interfaces, but also quantify them to investigate how the macrostructural dynamics of PFKL influences its behavior at a molecular level. This method can be used to probe the interactome of other proteins based on interacting interfaces under various conditions and further explore how form relates to function.

## Conclusion

PFKL self-assembles into activated filaments via interactions between interface 1 and 2 *in vitro*. It also self-assembles into inhibited puncta *in vivo*. The relation between these opposing modes of self-assembly was unclear, but we hypothesize inhibiting PFKL in cells would alter key protein-protein interactions with puncta. We used site-specific unnatural amino acid incorporation of a photochemical crosslinker AzF in combination with TMT labeled multiplexed proteomics to measure protein-protein interactions with interface 1 and 2 in PFKL. Our results revealed interface 2 as a hotspot of PFKL interactions with cytoskeletal and glycolytic proteins. Furthermore, we showed citrate inhibition attenuated PFKL interactions with interface 2, implying interface 2 was sequestered during puncta formation. These data suggest puncta structures contains PFKL filaments dependent on interface 2, which opposed the activated filament formation involving both interface 1 and interface 2.

## MATERIALS AND METHODS

### Plasmids, transfections, and antibodies

HEK293T cells were maintained in Dulbecco’s Modified Eagle’s Medium of high glucose with 10% fetal bovine serum, 1% penicillin/streptomycin, and 1% glutamine. Cells were transfected twenty-four hours after seeding 2×10^5^ cells/mL using the calcium phosphate transfection method. Approximately 3ug of AzF tRNA/RS DNA (AddGene Plasmid #105829) and 3ug of PFKL (tagged with the FLAG epitope) DNA were used for 10cm dishes. Around 18 hours of transfection, growth media was replaced with media containing 1mM of AzF (Chem Impex).

4ug of WT/mutant DNA plasmid, and 0.5ug of GFPTAG DNA plasmid were used. Lipofectamine 3000 was used to transfect Huh7 cells, and 3ug of tRNA/RS DNA plasmid, 3ug of WT/mutant DNA plasmid, and 0.5ug of GFPTAG DNA plasmid were used. Media was exchanged 18 hours after transfection, both for control samples and samples with unnatural amino acid, and cells were harvested 35-45 hours after transfection.

### Crosslinking and FLAG Immunoprecipitation

Before harvesting, AzF incorporation into PFKL mutants were confirmed by AcGFP1 expression. Cells were washed and resuspended in PBS and then exposed to 269 nm UV light for 1.5 minutes (1.1J total accumulated energy) to initiate photo-crosslinking. They were harvested immediately afterwards. FLAG immunoprecipitation was performed on the samples. Cells were lysed on the plate for 20 minutes at 4^°^C with radio-immunoprecipitation assay (RIPA) buffer (50mM Tris pH 7.5, 150mM NaCl, 0.1% SDS, 1% Triton X-100, 0.5% deoxycholate) containing Roche protease inhibitor. Centrifugation at 21.1g at 4°C cleared lysates. Pierce BCA Protein Assay Kit was used to normalize protein concentrations. Sigma Sepharose 4B and GenScript FLAG beads were washed three times with PBS. Normalized protein samples were added to sepharose beads and rocked at 4^°^C for one hour and then transferred to FLAG beads for overnight incubation on rocker at 4^°^C. The next day, beads were spun down at 400g for 3 minutes in a 4^°^C centrifuge. FLAG resin was washed four times with RIPA buffer and then eluted twice with 3x Laemmli buffer (6% SDS, 62.5mM Tris) for five minutes at 95^°^C. After elution, DTT was added to the samples at a final concentration of 17 mM and they were boiled once more for five minutes at 95^°^C before loading onto 10% SDS-PAGE gel that was run at 60V for 20 minutes and then 160V for one hour. Immunoprecipitation was confirmed via Western blot using 1:1000 dilutions of M2 anti-FLAG antibodies (Sigma-Aldrich, F1804).

### Multiplexed LC-MS/MS

The elution samples from the immunoprecipitations were prepared for mass spectrometry. A 3:1:3 ratio of methanol/chloroform/water was used to precipitate the eluted proteins, which were then washed twice with methanol. After each methanol wash, samples were spun down at 21,100 .1g for five minutes at room temperature. The methanol then evaporated by leaving samples exposed to air for 2-3 hours. The samples were then resuspended in 1% Waters Rapigest SF, reduced with 5 mM TCEP (Sigma), alkylated with 10 mM iodoacetamide (Sigma), and digested with 0.5 μg of trypsin/Lys-C, (Sequencing Grade, Promega, or Pierce) in 50mM HEPES (pH 8.0) at 37°C with 750 RPM shaking overnight. The following day, peptides were labelled using ThermoScientific 16-plex TMTpro for 1 hour and quenched with 10% w/v ammonium bicarbonate for 1 hour. The samples were then combined and acidified with ∼200 µL mass spec grade formic acid. After concentrating the sample on the rotary evaporator and resuspending in 1500 μL of buffer A (95% water, 4.9% acetonitrile, and 0.1% formic acid), the Rapigest products were then discarded by spinning sample down at 21.1g for 30 minutes.

### MudPIT LC-MS/MS Analysis

1.5cm C18 resin, 1.5cm SCX resin, and 1.5cm C18 resin were stacked in alternate layers to prepare triphasic MudPIT columns as previously stated in the literature^24^. TMT-labelled samples (20ug) were loaded onto the microcapillaries through a high-pressure chamber and a 30-minute wash in buffer A was subsequently conducted. The Ultimate 3000 nanoLC system was used for liquid chromatography and peptides were fractionated online. The Exploris 480 mass spectrometer (Thermo Fisher) was used for peptide analysis. The LC column switching valve had the MudPIT columns installed on it, followed by a 20cm fused silica microcapillary column containing Aqua C18, 3µm, C18 resin (Phenomenex) ending in a tip pulled by a laser. Before use, columns and MudPIT capillaries were both washed. Sequential injections (10µl) of 0%, 10%, 20%, 30%, 40%, 50%, 60%, 70%, 80%, 90%, and 100% of buffer C (500mM ammonium acetate, 94.9% water, 5% acetonitrile, 0.1% formic acid) and a subsequent injection of 90% buffer C and 10% buffer B (99.9% acetonitrile, 0.1% formic acid v/v) constituted the MudPIT runs. Using a flow rate of 500nL/min, a 130 minute gradient followed each injection (0-6 min: 2% buffer B, 8 min: 5% B, 100 min: 35% B, 105min: 65% B, 106-113 min: 85% B, 113-130 min: 2% B). Using a spray voltage of 2.2kV as well as an ion transfer tube temperature of 275°C and a RF Lens of 40%, electrospray ionization was performed directly from the tip of the microcapillary column. A scan range of 400-1600m/z, automatic injection times, an AGC target of 300%, and 120k resolution, MS1 spectra were collected. A charge state 2-7, TopSpeed method (3s cycle time), isolation window 0.4 m/z, HCD fragmentation using a normalized collision energy of 36%, resolution 45k, AGC target of 200%, automatic maximum injection times, and a dynamic exclusion (20 ppm window) set to 60s were used to collect data-dependent MS2 spectra using a monoisotopic peak selection mode: peptide. All proteins identified by condition are list in **Supplemental Table 5**.

### Crosslinked interaction identification and peptide identification and quantification

Crosslinked bands on the Western blot (indicating proteins that interact with PFKL) were identified as those with a higher molecular weight than the full-size WT and mutant PFKL with AzF incorporated (around 75 kDa). The full-length AcGFP1 PFKL mutants with AzF incorporated were around 100kDa. Band quantifications were performed using ImageLab. Thermo Fisher Proteome Discoverer 2.4 using SwissProt human database allowed for identification and quantification of proteins (other than WT PFKL and mutants) enriched in eluted samples. Peptide abundances were normalized according to the total amount of protein in each channel.

### PFK Enzyme Activity Assay

Sigma-Aldrich Phosphofructokinase (PFK) Activity Colorimetric Assay Kit was used to test the physiological activity of the PFKL mutants on whole cell lysates at dilution of 1:1000 in PFK assay buffer. Kinetic reaction assay was performed in a 96 well plate at room temperature for 1 hour, taking readings every minute and shaking 5 times between reads.

### Addition of Citrate and Imaging of PFKL Structures

To image PFKL structures in cells, HEK293T cells were transiently transfected using the lipofectamine3000 method with either WT FLAG-AcGFP1-PFKL or a TAG mutant (as indicated). TAG mutants were co-transfected with an AzF tRNA/RS construct. Transfections were performed at 80% confluency and cells were plated in 35 mm glass bottomed dishes (Ibidi #81218). Four to six hours after transfection, media was exchanged for DMEM supplemented with 10 % FBS 1% penicillin/streptomycin to remove traces of lipofectamine and trypsin. WT cells were imaged beginning at 18 hours after transfection. TAG mutant expressing cells were exchanged into media containing 1 mM AzF at 16 hours after transfection and imaged 18 hours after transfection. Before imaging, media was exchanged for FluoroBrite Dulbecco’s Modified Eagle’s Medium (Thermo Fisher) with 10% fetal bovine serum to avoid background absorbance by the phenol red present in Dulbecco’s Modified Eagle’s Medium. Cells were imaged before and after (10-minute incubation) addition of 20 mM sodium citrate.

### Statistical Analysis

To filter high-confidence interactors of PFKL proteins, we combined log2 fold enrichment and adjusted p-value calculated using a two tailed paired student t-test according to previously published approach (Keilhauer et al., 2015). Standard deviation (σ) of log2 protein abundance fold changes between PFKL transfected versus TD tomato mock transfected groups were calculated. 2σ or 1σ fold change cutoffs were used for high-confidence and medium-confidence interactors respectively as calculated with an adjusted p-values using hyperbolic curve y>c/(x−x0), where y is the adjusted p-value, x is the log2 fold change, x0 is the standard deviation (2σ or 1σ), and c is the curvature (c= 0.4 for 1σ and 0.8 for 2σ). Statistical significance between aggregated interacting proteins across pathways (Figure 5D) was calculated using a paired two tailed ratio student t test in Prism including all protein identifications with at least two replicates.

### Network Plots and Identification of Overlapping Secondary Interactors

Overlapping secondary interactors between the glycolysis, carbohydrate derivative metabolism, and cytoskeleton proteins identified were generated by scraping the STRING database using the python API. We first generated an extended secondary interactome by searching for the top 20 interactors using the limit parameter in STRING API and searching against the human proteome (species 9606). Then, we cross-compared the extended secondary interactome of each pathway with each other (e.g. glycolysis vs. carbohydrate derivative metabolism, glycolysis vs. cytoskeletal subunit etc.) an dropped any secondary interactors that did not appear in both pathways extended interactomes. Next, we concatenated the primary interactors from each pathway with overlapping secondary interactors into a single data set. Finally, the overlapping secondary interactors were searched in the STRING database human proteome to determine interactors between secondary interactors with a threshold of greater than 50% likelihood in the experimental score category. The results were plotted in Cytoscape.

### Fluorescence Microscopy

Imaging was performed on an epi-fluorescence microscope (Axio Observer 3, Zeiss, White Plains, NY) based on the inverted-type microscope, coupled to a white LED (X–Cite mini+, Excelitas, Waltham, MA) as an excitation light source. A 63 X objective was used (N-Achroplan, 63x/0.85 M27, Item no.: 420980-9900, Zeiss, Oberkochen, Germany). AcGFP1 was excited with blue light generated by using white light from the LED source passed through a ET480/40X bandpass filter (Chroma, Bellows Falls, VT) and a FF509-FDi01 dichroic mirror (Semrock, West Henrietta, NY). The emission was passed through an ET510lp filter (Chroma). Images were collected using 200 ms exposure time at 1×1 binning. Images were processed using FIJI (Schindelin et al., 2012).

## ACKNOWLEDGEMENTS

This work was supported by NIH grants R35 GM133552, R35 GM151146, The Research Corporation for the Advancement of Science, and the Gordon and Betty Moore Foundation. EFM was supported by a predoctoral fellowship from the National Heart, Lung, and Blood Institute (F31 HL162483-01A1). S.O.S. was partially supported by the National Science Foundation Graduate Research Fellowship under Grant No. DGE-2139841 and the National Institutes of Health under Biophysics Training Grant No. T32 GM 008283.

## AUTHOR CONTRIBUTIONS

Conceptualization, C.M.D., L.P. Funding acquisition, E.F.M., C.M.D., L.P. Data Collection, E.F.M., A.S., S.O.S., Data Analysis, E.F.M., A.S., S.O.S., Writing_original draft, A.S., E.F.M.; Writing_review and editing, E.F.M., A.S., C.M.D., and L.P. All authors have read and agreed to the published version of the manuscript.

## Notes

### Competing Interest Statement

The authors have declared no competing interest.

## REFERENCES

Akram, M., 2013. Mini-review on Glycolysis and Cancer. J. Cancer Educ. 28, 454–457. 10.1007/s13187-013-0486-9

Amara, N., Cooper, M.P., Voronkova, M.A., Webb, B.A., Lynch, E.M., Kollman, J.M., Ma, T., Yu, K., Lai, Z., Sangaraju, D., Kayagaki, N., Newton, K., Bogyo, M., Staben, S.T., Dixit, V.M., 2021. Selective activation of PFKL suppresses the phagocytic oxidative burst. Cell 184, 4480–4494.e15. 10.1016/j.cell.2021.07.004

Campanella, M.E., Chu, H., Low, P.S., 2005. Assembly and regulation of a glycolytic enzyme complex on the human erythrocyte membrane. Proc. Natl. Acad. Sci. 102, 2402–2407. 10.1073/pnas.0409741102

Chen, C., Peng, T., 2020. Protocol for Site-Specific Photo-Crosslinking Proteomics to Identify Protein-Protein Interactions in Mammalian Cells. STAR Protoc. 1, 100109. 10.1016/j.xpro.2020.100109

Chin, J.W., Cropp, T.A., Anderson, J.C., Mukherji, M., Zhang, Z., Schultz, P.G., 2003. An Expanded Eukaryotic Genetic Code. Science 301, 964–967. 10.1126/science.1084772

Davies, J.P., Almasy, K.M., McDonald, E.F., Plate, L., 2020. Comparative Multiplexed Interactomics of SARS-CoV-2 and Homologous Coronavirus Nonstructural Proteins Identifies Unique and Shared Host-Cell Dependencies. ACS Infect. Dis. 6, 3174–3189. 10.1021/acsinfecdis.0c00500

DeWane, G., Salvi, A.M., DeMali, K.A., 2021. Fueling the cytoskeleton – links between cell metabolism and actin remodeling. J. Cell Sci. 134, jcs248385. 10.1242/jcs.248385

El-Bacha, T., De Freitas, M.S., Sola-Penna, M., 2003. Cellular distribution of phosphofructokinase activity and implications to metabolic regulation in human breast cancer. Mol. Genet. Metab. 79, 294–299. 10.1016/S1096-7192(03)00117-3

Futran, A.S., Kyin, S., Shvartsman, S.Y., Link, A.J., 2015. Mapping the binding interface of ERK and transcriptional repressor Capicua using photocrosslinking. Proc. Natl. Acad. Sci. 112, 8590– 8595. 10.1073/pnas.1501373112

Gavin, A.-C., Bösche, M., Krause, R., Grandi, P., Marzioch, M., Bauer, A., Schultz, J., Rick, J.M., Michon, A.-M., Cruciat, C.-M., Remor, M., Höfert, C., Schelder, M., Brajenovic, M., Ruffner, H., Merino, A., Klein, K., Hudak, M., Dickson, D., Rudi, T., Gnau, V., Bauch, A., Bastuck, S., Huhse, B., Leutwein, C., Heurtier, M.-A., Copley, R.R., Edelmann, A., Querfurth, E., Rybin, V., Drewes, G., Raida, M., Bouwmeester, T., Bork, P., Seraphin, B., Kuster, B., Neubauer, G., Superti-Furga, G., 2002. Functional organization of the yeast proteome by systematic analysis of protein complexes. Nature 415, 141–147. 10.1038/415141a

Gomes Alves, G., Sola-Penna, M., 2003. Epinephrine modulates cellular distribution of muscle phosphofructokinase. Mol. Genet. Metab. 78, 302–306. 10.1016/S1096-7192(03)00037-4

Guo, X., Li, H., Xu, H., Woo, S., Dong, H., Lu, F., Lange, A.J., Wu, C., 2012. Glycolysis in the control of blood glucose homeostasis. Acta Pharm. Sin. B 2, 358–367. 10.1016/j.apsb.2012.06.002

Ha, C.E., Bhagavan, N.V., 2023. Carbohydrate metabolism I: glycolysis and the tricarboxylic acid cycle, in: Essentials of Medical Biochemistry. Elsevier, pp. 203–227. 10.1016/B978-0-323-88541-6.00030-2

Hashimoto, Y., Sheng, X., Murray-Nerger, L.A., Cristea, I.M., 2020. Temporal dynamics of protein complex formation and dissociation during human cytomegalovirus infection. Nat. Commun. 11, 806. 10.1038/s41467-020-14586-5

Hirschey, M.D., DeBerardinis, R.J., Diehl, A.M.E., Drew, J.E., Frezza, C., Green, M.F., Jones, L.W., Ko, Y.H., Le, A., Lea, M.A., Locasale, J.W., Longo, V.D., Lyssiotis, C.A., McDonnell, E., Mehrmohamadi, M., Michelotti, G., Muralidhar, V., Murphy, M.P., Pedersen, P.L., Poore, B., Raffaghello, L., Rathmell, J.C., Sivanand, S., Vander Heiden, M.G., Wellen, K.E., 2015. Dysregulated metabolism contributes to oncogenesis. Semin. Cancer Biol. 35, S129–S150. 10.1016/j.semcancer.2015.10.002

Ho, T., Potapenko, E., Davis, D.B., Merrins, M.J., 2023. A plasma membrane-associated glycolytic metabolon is functionally coupled to KATP channels in pancreatic α and β cells from humans and mice. Cell Rep. 42, 112394. 10.1016/j.celrep.2023.112394

Hunkeler, M., Hagmann, A., Stuttfeld, E., Chami, M., Guri, Y., Stahlberg, H., Maier, T., 2018. Structural basis for regulation of human acetyl-CoA carboxylase. Nature 558, 470–474. 10.1038/s41586-018-0201-4

Jang, S., Nelson, J.C., Bend, E.G., Rodríguez-Laureano, L., Tueros, F.G., Cartagenova, L., Underwood, K., Jorgensen, E.M., Colón-Ramos, D.A., 2016. Glycolytic Enzymes Localize to Synapses under Energy Stress to Support Synaptic Function. Neuron 90, 278–291. 10.1016/j.neuron.2016.03.011

Jang, S., Xuan, Z., Lagoy, R.C., Jawerth, L.M., Gonzalez, I.J., Singh, M., Prashad, S., Kim, H.S., Patel, A., Albrecht, D.R., Hyman, A.A., Colón-Ramos, D.A., 2021. Phosphofructokinase relocalizes into subcellular compartments with liquid-like properties in vivo. Biophys. J. 120, 1170–1186. 10.1016/j.bpj.2020.08.002

Kastritis, P.L., Gavin, A.-C., 2018. Enzymatic complexes across scales. Essays Biochem. 62, 501–514. 10.1042/EBC20180008

Keilhauer, E.C., Hein, M.Y., Mann, M., 2015. Accurate Protein Complex Retrieval by Affinity Enrichment Mass Spectrometry (AE-MS) Rather than Affinity Purification Mass Spectrometry (AP-MS). Mol. Cell. Proteomics 14, 120–135. 10.1074/mcp.M114.041012

Kemp, R.G., Foe, L.G., 1983. Allosteric regulatory properties of muscle phosphofructokinase. Mol. Cell. Biochem. 57, 147–154. 10.1007/BF00849191

Kemp, R.G., Gunasekera, D., 2002. Evolution of the Allosteric Ligand Sites of Mammalian Phosphofructo-1-kinase. Biochemistry 41, 9426–9430. 10.1021/bi020110d

Kim, M., McDonald, E.F., Sabusap, C.M.P., Timalsina, B., Joshi, D., Hong, J.S., Rab, A., Sorscher, E.J., Plate, L., 2023. Elexacaftor/VX-445-mediated CFTR interactome remodeling reveals differential correction driven by mutation-specific translational dynamics (preprint). Cell Biology. 10.1101/2023.02.04.527134

Kleiner, R.E., Hang, L.E., Molloy, K.R., Chait, B.T., Kapoor, T.M., 2018. A Chemical Proteomics Approach to Reveal Direct Protein-Protein Interactions in Living Cells. Cell Chem. Biol. 25, 110–120.e3. 10.1016/j.chembiol.2017.10.001

Kohnhorst, C.L., Kyoung, M., Jeon, M., Schmitt, D.L., Kennedy, E.L., Ramirez, J., Bracey, S.M., Luu, B.T., Russell, S.J., An, S., 2017. Identification of a multienzyme complex for glucose metabolism in living cells. J. Biol. Chem. 292, 9191–9203. 10.1074/jbc.M117.783050

Li, J., Cai, Z., Bomgarden, R.D., Pike, I., Kuhn, K., Rogers, J.C., Roberts, T.M., Gygi, S.P., Paulo, J.A., 2021. TMTpro-18plex: The Expanded and Complete Set of TMTpro Reagents for Sample Multiplexing. J. Proteome Res. 20, 2964–2972. 10.1021/acs.jproteome.1c00168

Lynch, E.M., Hicks, D.R., Shepherd, M., Endrizzi, J.A., Maker, A., Hansen, J.M., Barry, R.M., Gitai, Z., Baldwin, E.P., Kollman, J.M., 2017. Human CTP synthase filament structure reveals the active enzyme conformation. Nat. Struct. Mol. Biol. 24, 507–514. 10.1038/nsmb.3407

Makowski, M.M., Willems, E., Jansen, P.W.T.C., Vermeulen, M., 2016. Cross-linking immunoprecipitation-MS (xIP-MS): Topological Analysis of Chromatin-associated Protein Complexes Using Single Affinity Purification. Mol. Cell. Proteomics 15, 854–865. 10.1074/mcp.M115.053082

Masters, C., 1995. On the role of the cytoskeleton in metabolic compartmentation, in: The Cytoskeleton: A Multi-Volume Treatise. Elsevier, pp. 1–30. 10.1016/S1874-6020(06)80014-5

Masters, C., 1984. Interactions between glycolytic enzymes and components of the cytomatrix. J. Cell Biol. 99, 222s–225s. 10.1083/jcb.99.1.222s

Masters, C.J., Wilson, J.E., 1981. Interactions Between Soluble Enzymes and Subcellular Structur. Crit. Rev. Biochem. 11, 105–143. 10.3109/10409238109108700

McDonald, E.F., Sabusap, C.M.P., Kim, M., Plate, L., 2022. Distinct proteostasis states drive pharmacologic chaperone susceptibility for cystic fibrosis transmembrane conductance regulator misfolding mutants. Mol. Biol. Cell 33, ar62. 10.1091/mbc.E21-11-0578

Menard, L., Maughan, D., Vigoreaux, J., 2014. The Structural and Functional Coordination of Glycolytic Enzymes in Muscle: Evidence of a Metabolon? Biology 3, 623–644. 10.3390/biology3030623

Norris, V., Amar, P., Legent, G., Ripoll, C., Thellier, M., Ovádi, J., 2013. Sensor potency of the moonlighting enzyme-decorated cytoskeleton: the cytoskeleton as a metabolic sensor. BMC Biochem. 14, 3. 10.1186/1471-2091-14-3

Ovádi, J., Orosz, F., Hollán, S., 2004. Functional aspects of cellular microcompartmentation in the development of neurodegeneration: Mutation induced aberrant protein-protein associations. Mol. Cell. Biochem. 256, 83–93. 10.1023/B:MCBI.0000009860.86969.72

Ovádi, J., Saks, V., 2004. On the origin of intracellular compartmentation and organized metabolic systems. Mol. Cell. Biochem. 256, 5–12. 10.1023/B:MCBI.0000009855.14648.2c

Park, J.S., Burckhardt, C.J., Lazcano, R., Solis, L.M., Isogai, T., Li, L., Chen, C.S., Gao, B., Minna, J.D., Bachoo, R., DeBerardinis, R.J., Danuser, G., 2020. Mechanical regulation of glycolysis via cytoskeleton architecture. Nature 578, 621–626. 10.1038/s41586-020-1998-1

Plate, L., Rius, B., Nguyen, B., Genereux, J.C., Kelly, J.W., Wiseman, R.L., 2019. Quantitative Interactome Proteomics Reveals a Molecular Basis for ATF6-Dependent Regulation of a Destabilized Amyloidogenic Protein. Cell Chem. Biol. 26, 913–925.e4. 10.1016/j.chembiol.2019.04.001

Real-Hohn, A., Zancan, P., Da Silva, D., Martins, E.R., Salgado, L.T., Mermelstein, C.S., Gomes, A.M.O., Sola-Penna, M., 2010. Filamentous actin and its associated binding proteins are the stimulatory site for 6-phosphofructo-1-kinase association within the membrane of human erythrocytes. Biochimie 92, 538–544. 10.1016/j.biochi.2010.01.023

Richards, A.L., Eckhardt, M., Krogan, N.J., 2021. Mass spectrometry_-_based protein–protein interaction networks for the study of human diseases. Mol. Syst. Biol. 17, e8792. 10.15252/msb.20188792

Roberts, S.J., Somero, G.N., 1987. Binding of phosphofructokinase to filamentous actin. Biochemistry 26, 3437–3442.

Schindelin, J., Arganda-Carreras, I., Frise, E., Kaynig, V., Longair, M., Pietzsch, T., Preibisch, S., Rueden, C., Saalfeld, S., Schmid, B., Tinevez, J.-Y., White, D.J., Hartenstein, V., Eliceiri, K., Tomancak, P., Cardona, A., 2012. Fiji: an open-source platform for biological-image analysis. Nat. Methods 9, 676–682. 10.1038/nmeth.2019

Seidel, L., Zarzycka, B., Zaidi, S.A., Katritch, V., Coin, I., 2017. Structural insight into the activation of a class B G-protein-coupled receptor by peptide hormones in live human cells. eLife 6, e27711. 10.7554/eLife.27711

Shah, U.H., Toneatti, R., Gaitonde, S.A., Shin, J.M., González-Maeso, J., 2020. Site-Specific Incorporation of Genetically Encoded Photo-Crosslinkers Locates the Heteromeric Interface of a GPCR Complex in Living Cells. Cell Chem. Biol. 27, 1308–1317.e4. 10.1016/j.chembiol.2020.07.006

Stoddard, P.R., Lynch, E.M., Farrell, D.P., Dosey, A.M., DiMaio, F., Williams, T.A., Kollman, J.M., Murray, A.W., Garner, E.C., 2020. Polymerization in the actin ATPase clan regulates hexokinase activity in yeast. Science 367, 1039–1042. 10.1126/science.aay5359

Szklarczyk, D., Gable, A.L., Lyon, D., Junge, A., Wyder, S., Huerta-Cepas, J., Simonovic, M., Doncheva, N.T., Morris, J.H., Bork, P., Jensen, L.J., Mering, C. von, 2019. STRING v11: protein–protein association networks with increased coverage, supporting functional discovery in genome-wide experimental datasets. Nucleic Acids Res. 47, D607–D613. 10.1093/nar/gky1131

Usenik, A., Legiša, M., 2010. Evolution of Allosteric Citrate Binding Sites on 6-phosphofructo-1-kinase. PLoS ONE 5, e15447. 10.1371/journal.pone.0015447

Webb, B.A., Dosey, A.M., Wittmann, T., Kollman, J.M., Barber, D.L., 2017. The glycolytic enzyme phosphofructokinase-1 assembles into filaments. J. Cell Biol. 216, 2305–2313. 10.1083/jcb.201701084

Wright, M.T., Kouba, L., Plate, L., 2021. Thyroglobulin Interactome Profiling Defines Altered Proteostasis Topology Associated With Thyroid Dyshormonogenesis. Mol. Cell. Proteomics 20, 100008. 10.1074/mcp.RA120.002168

Wu, X., Spence, J.S., Das, T., Yuan, X., Chen, C., Zhang, Y., Li, Y., Sun, Y., Chandran, K., Hang, H.C., Peng, T., 2020. Site-Specific Photo-Crosslinking Proteomics Reveal Regulation of IFITM3 Trafficking and Turnover by VCP/p97 ATPase. Cell Chem. Biol. 27, 571–585.e6. 10.1016/j.chembiol.2020.03.004

Zhang, W.H., Otting, G., Jackson, C.J., 2013. Protein engineering with unnatural amino acids. Curr. Opin. Struct. Biol. 23, 581–587. 10.1016/j.sbi.2013.06.009

