## Supplemental Information for "Site-Specific Crosslinking Reveals Phosphofructokinase-L Inhibition Drives Self-Assembly and Attenuation of Protein Interactions"

Table of Contents

Supplemental Figures……………………………………………………………………………………..2-6

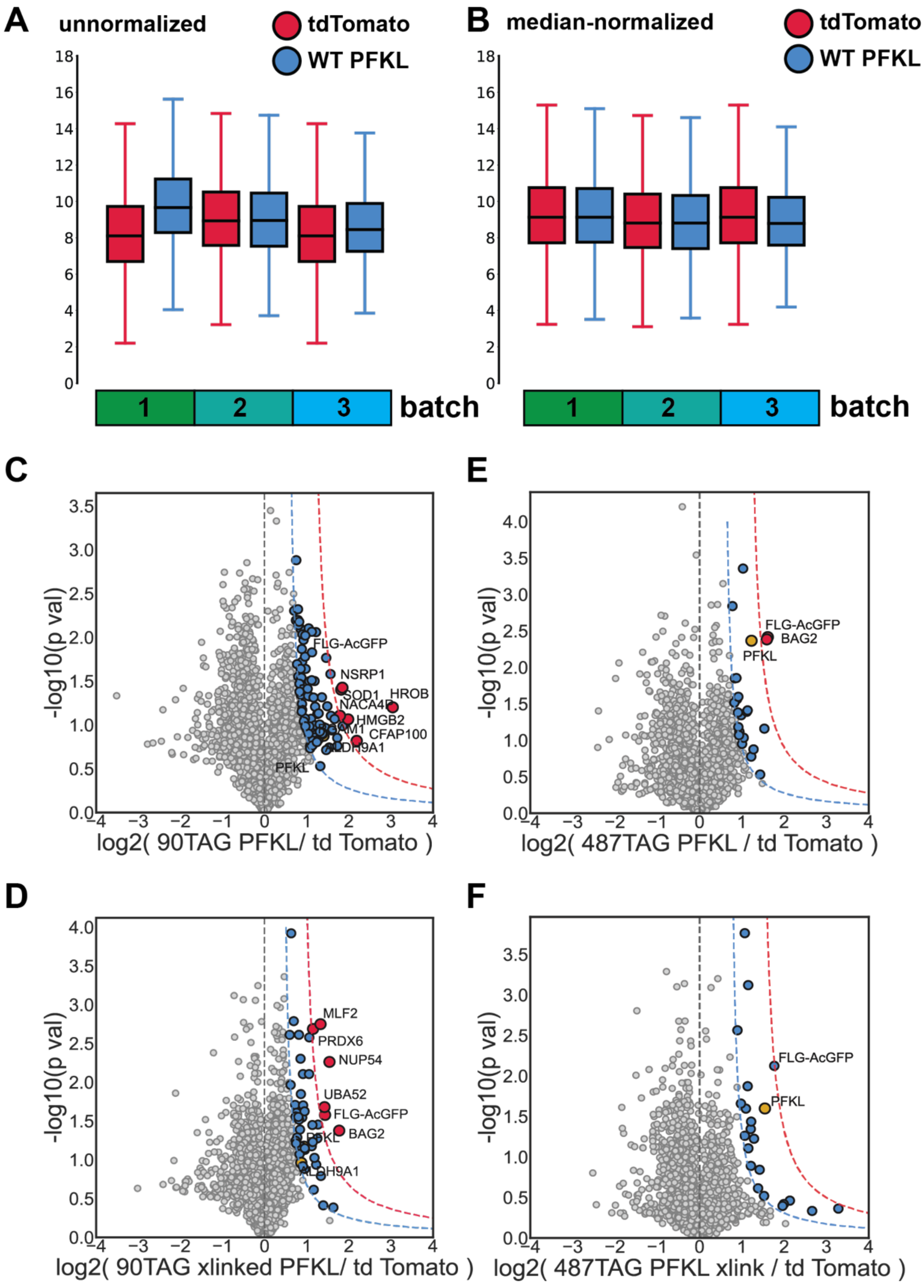


**Supplemental Figure S1. Median normalization of TMT channels and identification of K90AzF and Y487AzF interactors.** (**A, B**) Box and whisker plot showing the TMT intensity distributions of tdTomato and WT PFKL for all 3 LC-MS/MS runs conducted in this study. (**A**) The sum of all TMT intensities in the tdTomato and WT PFKL channels were normalized to total peptide level. (**B)** To make the median in each run for all samples equal, the sum of all TMT intensities in each channel were normalized across runs, as represented by tdTomato and WT PFKL here. Intensities within a given batch are more closely aligned. (**C, D, E, F)** Volcano plots of K90AzF (**C**), K90AzF exposed to

UV light (**D**), Y487AzF (**E**), and Y487AzF exposed to UV light (**F**) for identification of interactors. The x-axis represents the log2 fold change of K90AzF (**C**,**D**) or Y487AzF (**E**,**F**) TMT quantification compared to the tdTomato control TMT quantification for three biological replicates. The y-axis represents the -log10 transformed p-values, as calculated by student's t-test. Bait protein is labeled as PFKL and FLG-AcGFP. High-confidence (red) and medium-confidence (blue) interactors are demarcated at 1σ' and 2 σ' respectively. Glycolytic interactors are represented in yellow. K90AzF, located in the active site, demonstrates a high number of interactors (**C**,**D**), while only one interactor was found to be enriched with Y487TAG, located in interface 1 (**E**,**F**).

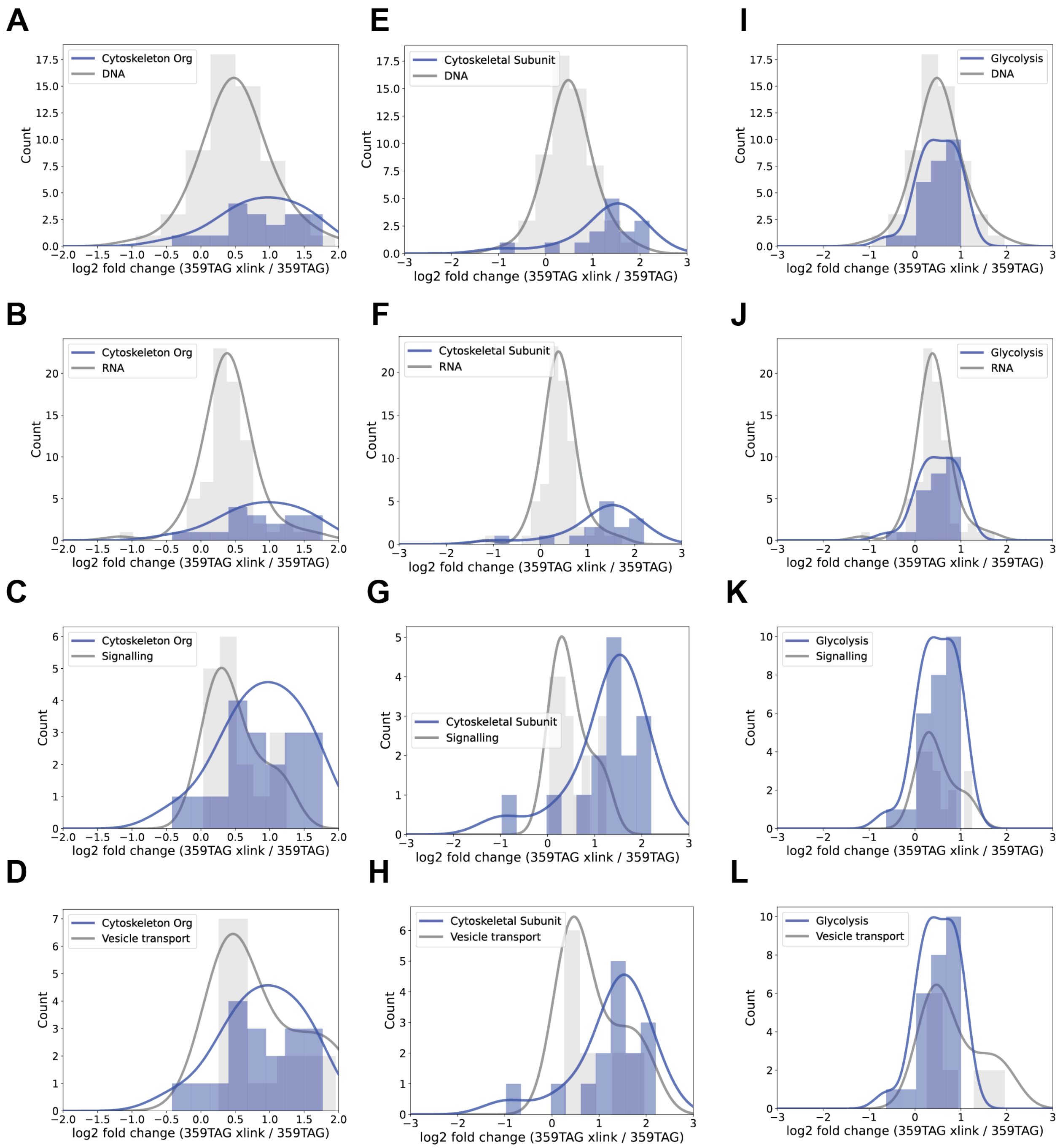


**Supplemental Figure S2. Comparison of interactors in various cellular pathways.** Histograms displaying the log2 fold change of Q359AzF interactors functioning in different pathways, including cystokeleton organization, DNA, RNA, glycolysis, signaling, and vesicle transport, to access the processes which are most represented in the Q359AzF interactome. Within each histogram, the log2 fold change distribution for each pathway is overlaid in gray and purple for comparison. All curves are fitted to the Gaussian distribution with 0.6 as the covariance factor. Compared to other pathways, proteins functioning in cytoskeleton organization are generally more enriched as interactors of Q359AzF (**A**-**H**). Glycolytic proteins are less enriched compared to DNA and RNA pathways (**I**,**J**) but more prevalent compared to signaling and vesicle transport pathways (**K**,**L**).

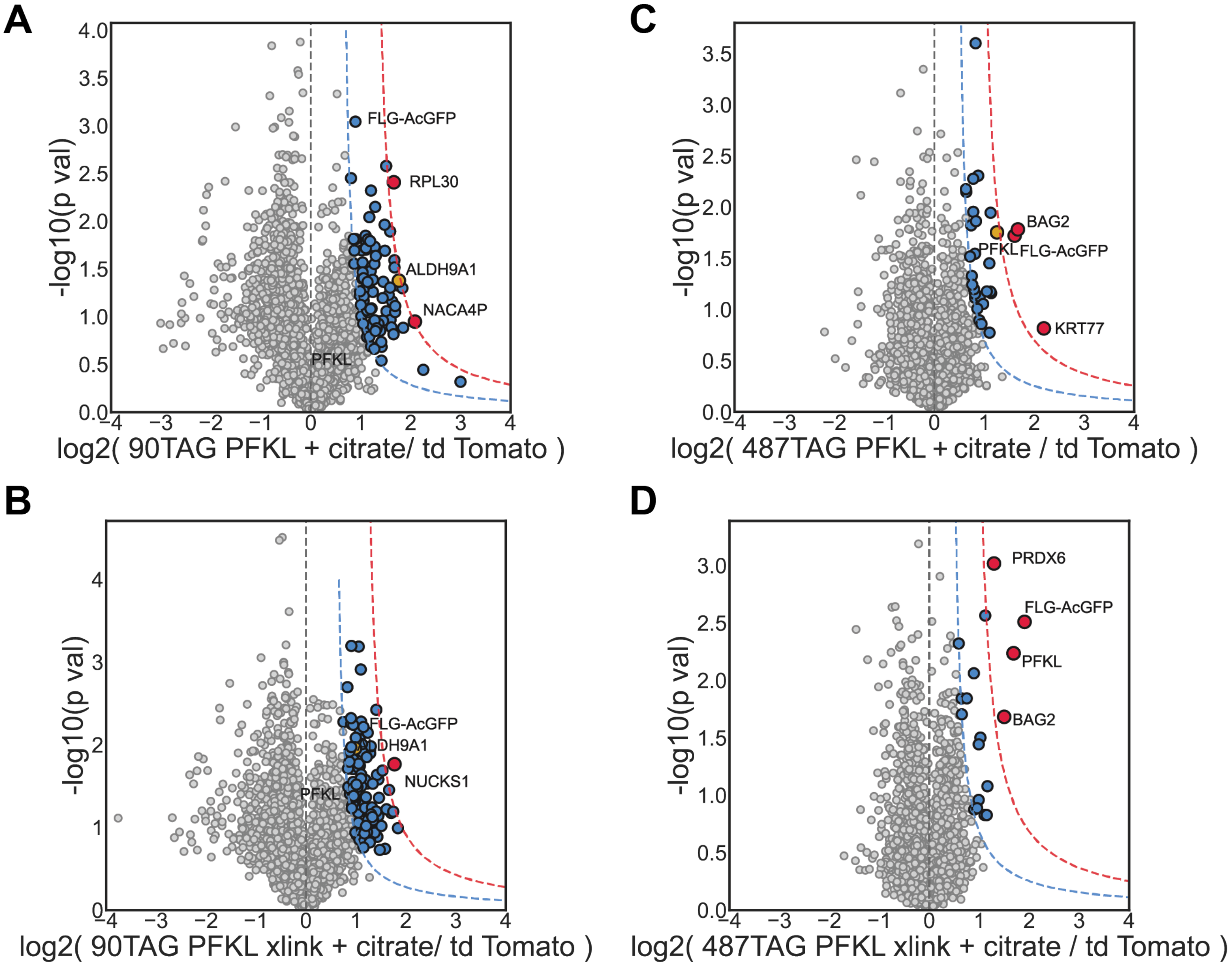


**Supplemental Figure S3. Quantification and identification of K90AzF and Y487AzF interactors under citrate-induced enzymatic inhibition.** Volcano plots of K90AzF with citrate (A), K90AzF with citrate and exposed to UV light (B), Y487AzF with citrate (C), and Y487AzF with citrate and exposed to UV light (D) for identification of interactors. The X axis represents the log2 fold change of K90AzF (A,B) or Y487AzF (C,D) TMT quantification over the tdTomato TMT quantification across three biological replicates. The Y axis represents the -log10 transformed p-values, as calculated by student's t-test. Bait protein is labeled as PFKL and FLG-AcGFP. High-confidence (red) and medium-confidence (blue) interactors are demarcated at 1σ' and 2 σ' respectively. Glycolytic interactors are represented in yellow. Citrate slightly alters enzyme interactions in a qualitative manner, but the number of interactors stays approximately the same for both PFKL mutants.

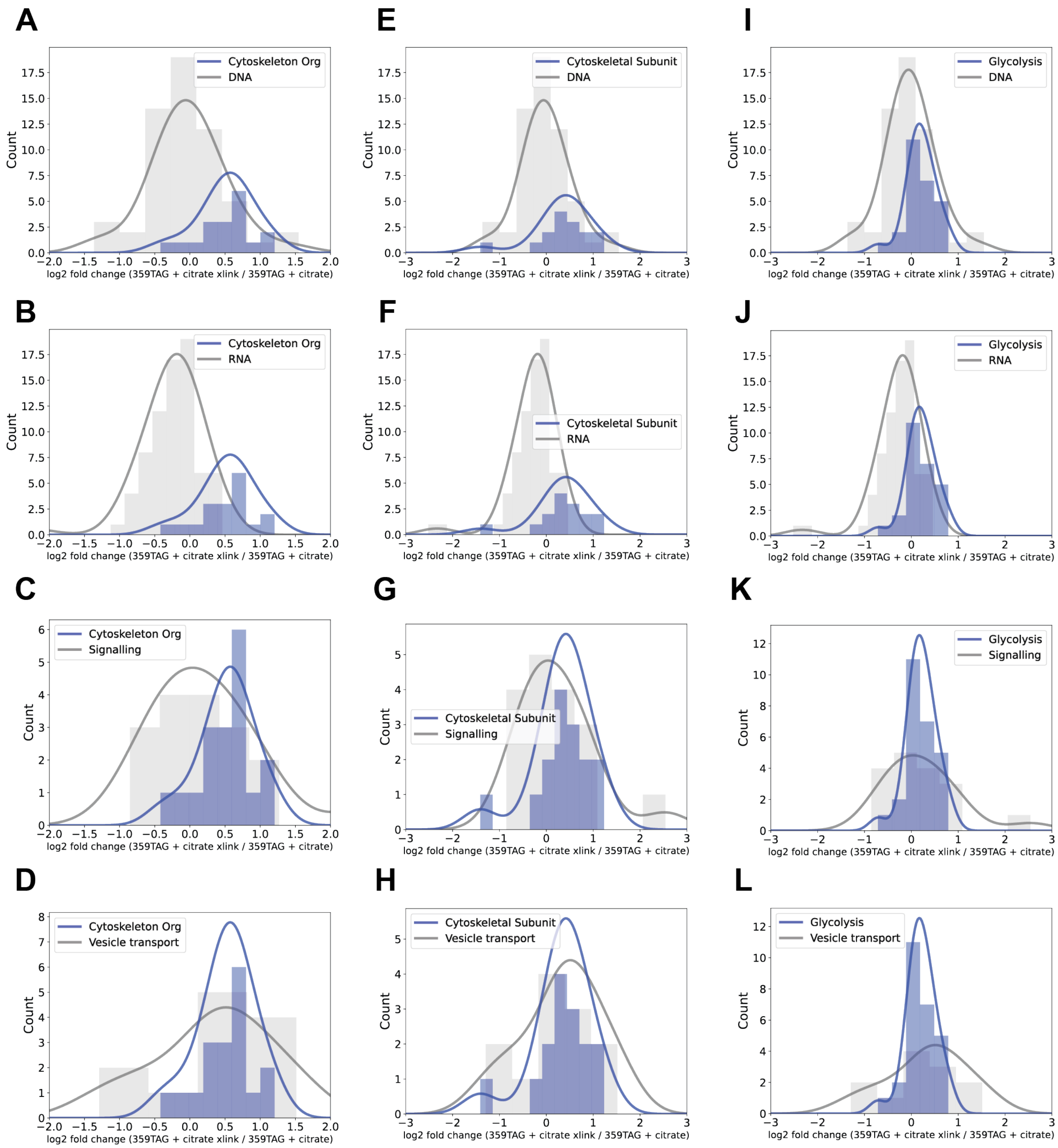


**Supplemental Figure S4. Comparison of interaction changes of Q359AzF PFKL under citrate treatment with various cellular pathways.** Histograms displaying the log2 fold change of Q359AzF interactors treated with citrate comparing the UV cross-linked to no cross-linking conditions. Enrichment of different pathways is shown, including cystokeleton organization, DNA, RNA, glycolysis, signaling, and vesicle transport, to access the processes which are most represented in the Q359AzF interactome. Within each histogram, the log2 fold change distribution for each pathway is overlaid in gray and purple for comparison. All curves are fitted to the Gaussian distribution with 0.6 as the covariance factor. Compared to other pathways, proteins functioning in cytoskeleton organization are generally more enriched as interactors of Q359AzF (A-H). Glycolytic proteins are less enriched compared to DNA and RNA pathways (I,J) but more prevalent compared to signaling and vesicle transport pathways (K,L).
